# Systematic evaluation of genome-wide metabolic landscapes in lactic acid bacteria reveals diet-induced and strain-specific probiotic idiosyncrasies

**DOI:** 10.1101/2021.06.20.449192

**Authors:** Lokanand Koduru, Meiyappan Lakshmanan, Pei-Yu Lim, Pooi-Leng Ho, Mazlina Banu, Doo-Sang Park, Dave Siak-Wei Ow, Dong-Yup Lee

## Abstract

Lactic acid bacteria (LAB) naturally occur in animal and plant niches and are well-known to elicit several health benefits in humans. Yet, how they adapt their functional metabolic landscapes to diverse nutrient environments and synthesize relevant bioactive compounds remain unexplored across genera, species and strains. Hence, presented herein is a systematic framework for comprehensively characterizing the genome-wide metabolisms of six representative LAB by combining multi-omics data with *in silico* modeling. We analyse the differences in their growth and cellular fitness, biosynthetic capability of health-relevant compounds, i.e., postbiotics, and probable interactions with 15 common gut microbiota under 11 virtual dietary regimes, and show such attributes are diet- and species-specific. Particularly, some LAB exhibit a desirable balance between synthesis of beneficial postbiotic compounds, positive interactions with beneficial gut commensals, and the ability to colonize and persist in gut environment. We also observe that “high fat-low carb” diets likely lead to detrimental outcomes in most LAB. Our results clearly highlight that probiotics are not “one size fits all” commodities and need to be formulated in a personalised manner for their use as dietary supplements and live biotherapeutics. Overall, the proposed framework will systematize the probiotic administration and could also widen the strain repertoire.

---

‘Lactic acid bacteria (LAB)’ refers to a group of microaerophilic, Gram-positive bacteria which primarily ferment the hexose sugars into lactate^1^. LAB include several genera, *Enterococcus, Lactobacillus, Lactococcus, Leuconostoc, Oenococcus, Pediococcus* and *Streptococcus*, with functional classification into three major fermentative groups: homolactic, facultative heterolactic and obligative heterolactic, depending on the fermentation products which result from carbohydrate catabolism^2^. LAB are indigenous species in human food habitat and are ubiquitously found in various nutrient-rich niches such as milk environments, vegetables and meats^3^. Moreover, they also constitute a part of the human microbiome in the large and small intestines and colon mucosal layers, exhibiting complex molecular cross-talk with the host and other microbiota to confer several beneficial health effects, e.g., antibacterial^4–7^, immunomodulatory and cytokine stimulatory^8,9^, free radical scavenging^10,11^ and antitumor activities^12,13^. LAB have thus been a popular choice to maintain a good gut homeostasis and/or to ameliorate microbiota dysbiosis as it has been comprehensively reviewed elsewhere^14,15^.

Although certain LAB such as *Lactobacillus plantarum, Lactobacillus salivarius* and *Lactobacillus casei* are generally used as probiotics, their beneficial effects could be highly variable across different species, genera and strains. While several studies have extensively demonstrated the variations via both *in vitro* and *in vivo* experiments^16–23^, some have also reported a few commensurate health-promoting effects across various strains or even species^24–27^. Such conflicting observations possibly stem from a myriad of factors including the differences in evaluation methodology and the distinct mechanisms of action: while some beneficial effects, e.g. bile resistance^28^ or mucosal adhesion^29,30^, could be common across even genera^31^, others might be niche- or strain-specific, e.g. synthesis of bioactive compounds, i.e. postbiotics. Therefore, it is of high importance to comprehensively examine the unique metabolic capabilities of LAB and uncover their nutritional preferences and biosynthetic functions of various postbiotics in a context-specific manner. In this regard, the current availability of complete genome sequences for more than 1000 LAB strains^32,33^ has facilitated only a handful of comparative genomic analyses hinting at the plausible diversity in nutrient uptake and the ability to produce various antimicrobial substances^34,35^. Here, for the first time, we systematically evaluate the genome-wide functional metabolic capabilities of six representative LAB from different genera, species and fermentative groups, through an integrative framework based on comparative genomics, transcriptomics and *in silico* modelling, together with their growth and biochemical profiles obtained from a newly formulated chemically defined medium (CDM). Such comparative analyses allow us to elucidate the probiotic determinants and probable interactions of LAB with common gut commensal and pathogenic microbes under various dietary regimes.

## Results

### A common chemically defined medium for the unbiased evaluation of LAB growth characteristics

We selected six representative strains, *L. plantarum* WCFS1 (LbPt), *L. casei* subsp. *casei* ATCC 393 (LbCs), *Lactobacillus salivarius* ATCC 11741 (LbSv), *Lactobacillus fermentum* ATCC 14931 (LbFm), *Lactococcus lactis* subsp. *cremoris* NZ9000 (LcLt) and *Leuconostoc mesenteroides* subsp. *mesenteroides* ATCC 8293 (LeMt), from three different genera in order to cover a wide range of LAB across different species as well as diverse functional groups. Note that although many *Lactobacillus* species are well established as probiotics, here, we purposely chose representatives from other well-studied genera of LAB, *Leuconostoc* and *Lactococcus*, to provide a good balance in terms of species diversity. Following the selection of strains, we newly formulated a chemically defined medium which supports the growth of all LAB tested. Design of CDM is highly necessary to enable an unbiased and comprehensive metabolic characterisation of all six LAB strains which are generally auxotrophic to several nutrients, particularly amino acids and vitamins. Here, it should be emphasized that although rich nutrient media such as M17 or *Lactobacillus* broth might support their growth, the chemically undefined components in the media may introduce batch-batch variations and uncertainties to substrate/product measurements. We initially cultivated all six LAB in a CDM which was originally developed to grow *Lactococcus* and *Streptococcus* species^36^. However, this medium was only able to support the growth of LcLt and LbSv. Therefore, we modified it iteratively by including several key metabolites based on the known growth requirements of each LAB, hereafter referred as LAB defined medium (LABDM), for their unrestricted growth. Importantly, we eliminated the need for preparing multiple micronutrient component stocks of vitamins and minor salts in LABDM by replacing them with yeast nitrogen base (YNB), which is a defined commercial formulation of micronutrients widely used for yeast cultivation. The optimal supplementation of YNB ensured proper growth with minimum possible lag phase for most LAB strains (see **Methods** and **Supplementary Dataset S1**).

We next cultivated the LAB strains in LABDM and evaluated their batch growth characteristics under anaerobic conditions. Of them, LbPt and LbSv grew much faster, i.e. smaller doubling time than others (**Figure 1a**; p-val = 0.0077, Kruskal-Wallis test). However, LbSv exhibited significantly longer lag phase compared to other LAB (**Figure 1b and Supplementary Figure S1**; p-val = 0.0083, Kruskal-Wallis test). The concentrations of major nutrients such as glucose and amino acids, and primary by-product, lactate, in the LABDM were also monitored over the exponential phase. No significant difference was observed in the biomass yield from glucose across LAB although obligate heterofermentative LAB (LeMt and LbFm) showed slightly lower yields (**Figure 1c**; p-val = 0.0186, Kruskal-Wallis test), indicating their limited ability to catabolize amino acids and other nutrients for energy production, concordant with our previous study^37^.

**Figure 1.**
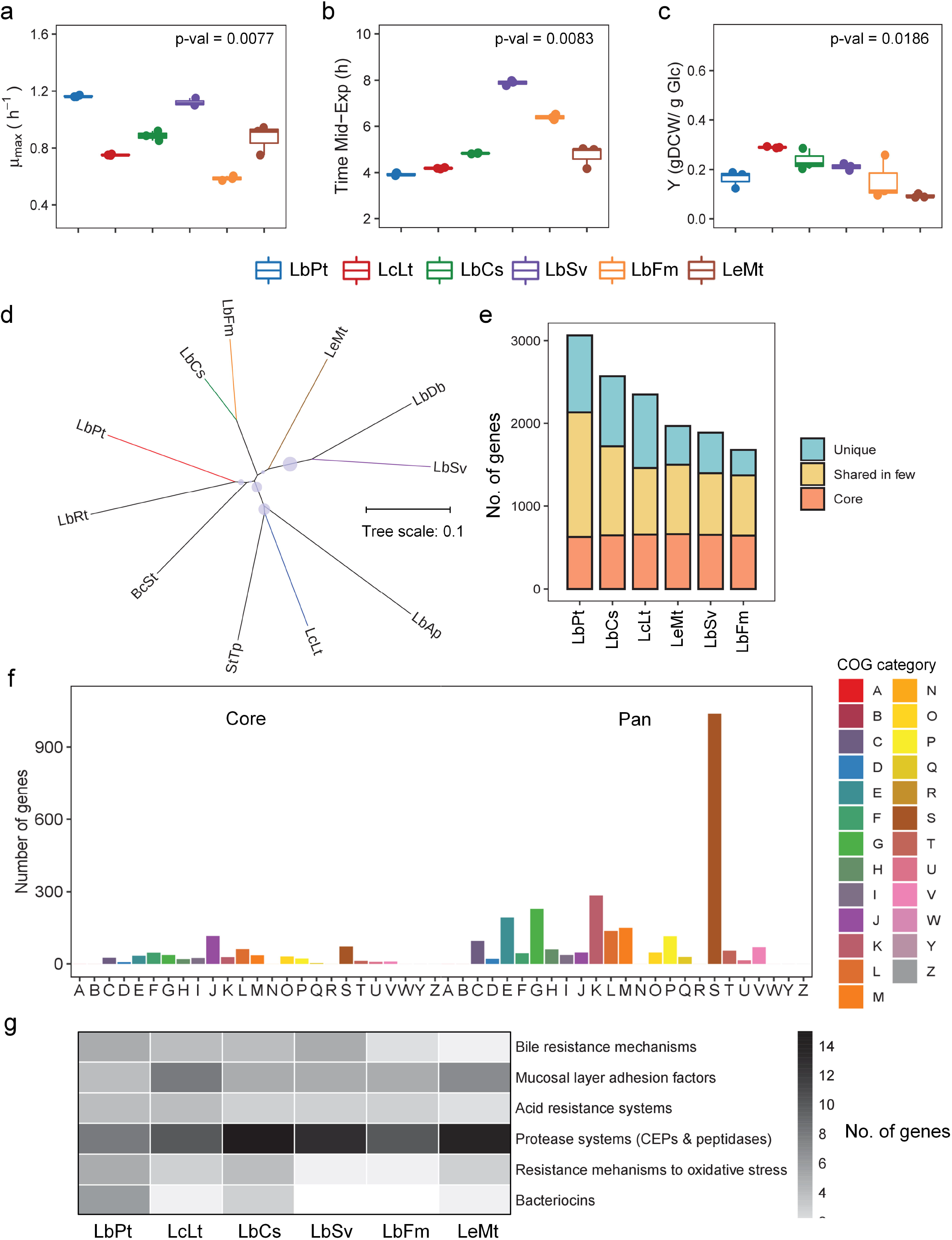
Growth phenotype and genomic characteristics of LAB strains. (a) Differences in the maximum growth rates across LAB in LABDM cultures (p-val = 0.0077, Kruskal-Wallis test), (b) differences in time required to reach mid-exponential phase across LAB in LABDM cultures (p-val = 0.0083, Kruskal-Wallis test), (c) Differences in yield of biomass over glucose across LAB in LABDM cultures (p-val = 0.0186, Kruskal-Wallis test), (d) phylogenetic tree showing the relationship between several LAB species with an outgroup member *B. subtilis* (BaSt), (e) the number of core and pan genes in LAB strains, (f) the functional annotation of core and pan genes using NOG mappings, (g) comparison of major non-metabolic genetic determinants influencing probiotic efficacy of LAB. Boxes in (a-c) include 1st to 3rd quartile of each population while whiskers extend up to 1.5 times the interquartile range from the closest box boundary. The size of circles in (d) represent the bootstrap values. Core genes in (f) are the genes which is present in all 6 LAB. Pan genes include both unique genes and the ones which ortholog only in a few. NOG categories in (f) is as follows: A – RNA processing and modification, B – Chromatin structure and dynamics, C – Energy production and conversion, D – Cell cycle control, cell division, chromosome partitioning, E – Amino acid transport and metabolism, F – Nucleotide transport and metabolism, G – Carbohydrate transport and metabolism, H – Coenzyme transport and metabolism, I – Lipid transport and metabolism, J – Translation, ribosomal structure and biogenesis, K – Transcription, L – Replication, recombination and repair, M – Cell wall/membrane/envelope biogenesis, N – Cell motility, O – Posttranslational modification, protein turnover, chaperones, P – Inorganic ion transport and metabolism, Q – Secondary metabolites biosynthesis, transport and catabolism, R – General function prediction only, S – Function unknown, T – Signal transduction mechanisms, U – Intracellular trafficking, secretion, and vesicular transport, V – Defense mechanisms, W – Extracellular structures, Y – Nuclear structure and Z – Cytoskeleton.

### Comparative genomic analysis of LAB reveals their pan genomic basis of probiotic capabilities

To understand how the probiotic characteristics vary across LAB, we conducted a thorough comparative genomic survey. We first constructed a phylogenetic tree using the genome sequences of all six species considered here with four other LAB and an outgroup member, *Bacillus subtilis* (BcSt) (see **Methods**). While most *Lactobacillus* strains cluster together, interestingly, LeMt and BcSt are also placed in the same cluster (**Figure 1d**). This confirms a previous observation that *Lactobacillus* genus is paraphyletic^34^ and underlines the remarkable genetic diversity of LAB species, which could be a result of their adaptation to various nutrient-rich niches. Furthermore, LAB genomes vary widely in size (2.55 ± 0.84 Mb), where LbPt and LbFm have the largest (3.84 Mb) and smallest (1.86 Mb) genomes, respectively (**Supplementary Table S1**), signifying the potential variations in probiotic traits among them. Such large variations in the genome size could possibly be a result of significant horizontal gene loss/gain in the evolutionary process, further confirming the ongoing process of genome degeneration as indicated by the presence of pseudogenes in most of the LAB^34^.

We next quantified the genetic redundancy across LAB on the basis of orthologous genes to comprehensively characterize how their genomic catalogues differ from each other (see **Methods**). A total of 627 unique “core” genes to be present commonly and 1567 “pan” genes that are either unique to a particular LAB or shared in a few were identified, thus emphasizing the extraordinary genetic diversity in LAB despite their small genome sizes (**Figure 1e**). Functional annotation of the core and pan genome (see **Methods**) highlighted that housekeeping processes such as translation, ribosomal structure and biogenesis, replication (17% compared to 1%) and post-translational modification, protein turnover, chaperones (5% compared to 2%) are highly enriched in the core genome compared to the pan genome. In addition, a few metabolic pathways such as lipid and nucleotide metabolisms, were highly conserved across LAB species (**Figure 1f**). On the other hand, more enriched in pan genome include carbohydrate, amino acid, inorganic ion, coenzyme and other secondary metabolism, and some non-metabolic categories such as transcription and function unknown. Significant enrichment of metabolic genes in pan genome, particularly carbohydrate and amino acid metabolisms, pinpoints the differences in their preferential adaptation to nutrient environments such as milk, plant and animals^38–40^ as well as the variations in their probiotic capabilities^41^. Moreover, the high proportion of genes with unknown function (980/3245) in LAB pan genome hints at the gaps in the current understanding and the need for a comprehensive functional annotation of such genes. LAB also synthesize a variety of proteins for resisting acid, oxidative and bile stresses, conferring anti-microbial activities, protein lysing proteases and anchor cells on host mucosal layer surfaces, which constitute important probiotic features. We mined all six LAB genomes and observed that LAB with larger genomes, i.e. LbPt and LbCs, synthesize multiple cell envelope proteinases (CEPs) and peptidases (**Figure 1g**; see **Methods** and **Supplementary Results**). Taken together, these findings suggest the dominant role of pan genomic features in determining the probiotic capabilities in LAB.

### Genome-wide transcriptome sequencing of LAB reveals unique gene expression profiles irrespective of their taxonomic groups

In order to delve deeper insights into the genome-wide transcriptional regulation among various taxonomic groups, we collected RNA at the mid-exponential phase from all strains and performed RNA-seq. At the outset, we compared the global gene expression profiles by analysing the distribution of genes which fall under various expression ranges. LbSv and LbFm have large number of genes in low expression range while LbPt, LbCs and LeMt have gene expression distribution slightly skewed towards high expression range (**Figure 2a** and **Supplementary Figure S2**). Subsequent enrichment analysis of NOG terms in these gene expression categories (see **Methods**) identified housekeeping functions, i.e. translation, ribosomal structure and biogenesis (FDR adj. p-val = 8.830 × 10^-30^ - 1.805 × 10^-17^, Fisher’s exact test), in high expression range (**Figure 2b**). We also noted that carbohydrate transport and metabolism is significantly enriched (FDR adj. p-val = 1.341 × 10^-10^ - 3.128 × 10^-7^, Fisher’s exact test) in low expression range particularly for LbPt, LbCs and LcLt, indicating that the metabolism of LAB with larger genomes is further regulated at the transcriptional level. Noticeably, we also found a wide spread expression of genes with unknown function, highlighting the need to further characterize their role in governing LAB phenotype.

**Figure 2.**
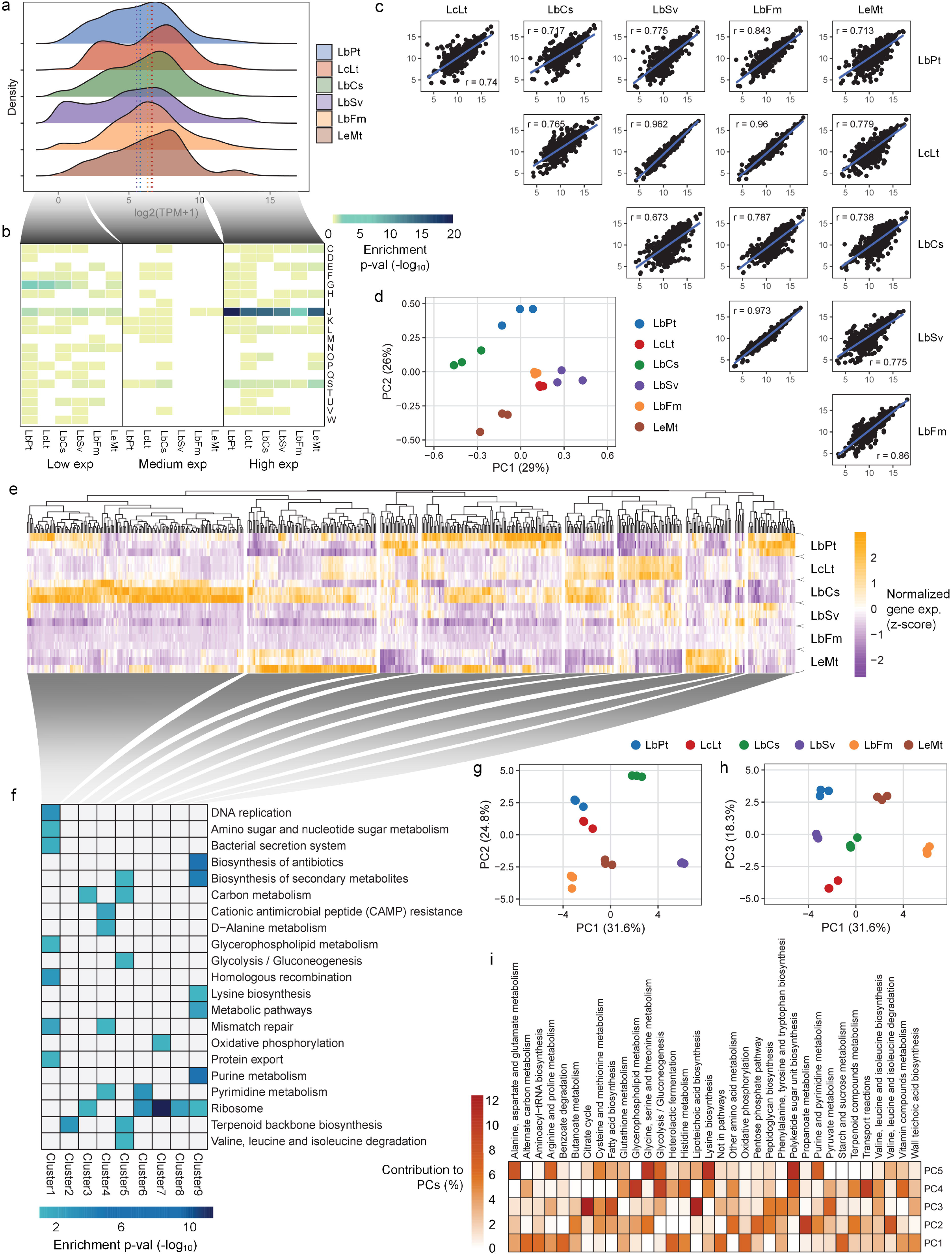
Transcriptomic characteristics of exponentially growing LAB in LABDM. (a) Cumulative distribution of gene expression values, (b) enrichment of NOG categories in three distinct gene expression ranges, i.e. low, medium and high, (c) plots showing pairwise correlation of gene expression of orthologous genes across various LAB, (d) principal component analysis (PCA) of the orthologous gene expression, (e) Hierarchical clustering of differentially expressed orthologous genes, (f) enrichment of KEGG pathways in differentially expressed genes and (g – j) PCA of metabolic genes in various LAB strains. Numbers in (c) represent the Spearman correlation coefficient, *r*. The NOG term abbreviations in (b) are same as **Figure 1**. Negative log10 transformations of enrichment p-value is provided in (b) and (f). Clustering was done with Euclidean distances of normalized gene expressions (Z-scores) across LAB in (e). PCA of metabolic genes was performed based gene assignments to various pathways in newly reconstructed GEMs.

In addition to the global transcriptome analysis, we compared the interspecies transcriptomic landscapes and their predisposition to various metabolic and probiotic capabilities by first identifying the 549 one-one orthologues (including 195 metabolic genes), and examining their normalized expression values using a set of housekeeping genes to correct for the species-specific bias (see **Methods** and **Supplementary Figure S3**). The expression profiles of these genes were analysed using two individual and complementary metrics: i) principal component analysis (PCA) and ii) correlation in a pairwise manner. Surprisingly, the gene expression patterns are markedly different between phylogenetically closer species such as LbSv and LbCs (Spearman correlation, *r* = 0.673) while it is highly similar in distant species (**Figures 2c-d**), e.g. LbFm and LcLt (Spearman correlation, *r* = 0.96). Hierarchical clustering of differentially expressed orthologous genes resulted in 9 major groups where each cluster is mainly represented by genes that are upregulated in a particular LAB (**Figure 2e**). Enrichment of functional categories in these genes further revealed that the carbon metabolism and several growth-related biological processes such as DNA replication (FDR adj. p-val = 3.162 × 10^-17^, modified Fisher’s exact test), ribosome and DNA mismatch repair (FDR adj. p-val = 0.0031, modified Fisher’s exact test) were upregulated in LAB with larger genomes such as LbPt, LbCs and LcLt, thus providing further evidences to why these species grow much faster than others (**Figure 2f** and **Supplementary Figure S4**). On the other hand, LeMt show high gene expression of anaerobic respiration (FDR adj. p-val = 0.0034, Fisher’s exact test), i.e. F1ATPase and menaquinone biosynthesis, indicating that redox-controlled metabolism governs the fastidious growth as reported earlier^37^. We also observed a high expression of certain postbiotics biosynthetic pathways in LbCs and LbPt, while it is lower in LbFm and LbSv.

Since the transcriptional regulation of each LAB is significantly different even among the phylogenetically closer species, we further performed a PCA focusing on metabolic gene expression. The first and second PCs account for ~32% and 25% variations, respectively, with major contributions from the functional categories relevant to growth (alternate carbon metabolism, aminoacyl-tRNA biosynthesis, heterolactic fermentation, oxidative phosphorylation and starch and sucrose metabolism) and biosynthesis of bioactive compounds (protonate metabolism, butanoate metabolism, lipoteichoic acid biosynthesis and vitamin metabolism) (**Figure 2g-i**). These results clearly suggest that gene expression variations among LAB can be better captured by their ability to adapt to various environments and produce relevant bioactive compounds rather than their taxonomic differences.

### Genome-scale modeling delineates the diverse metabolic repertoire and the predominant use of low energy-yielding pathways in LAB strains

As the comparative genomics and transcriptomics analyses revealed an unprecedented divergence in metabolic and physiological capabilities of LAB, we reconstructed new genomescale metabolic models (GEMs) for LbCs, LbFm and LbSv and updated the existing ones for LbPt^42^, LcLt^43^ and LeMt^37^, following the standard procedure^44^ (see **Methods**) to further probe their functional metabolic states. The resulting GEMs contained 686 ± 139 genes, 1042 ± 65 reactions and 906 ± 29 unique metabolites on an average, where LbFm and LbPt represented the smallest and largest models, respectively (**Supplementary Table S2**). We subsequently calculated the metabolic distances between pairs of any two GEMs using their genome-wide reactome and observed that all *Lactobacillus* strains shared more reactions than taxonomically distant LeMt and LcLt (**Supplementary Figure S5**). We then compared the LAB GEMs to identify the “core” and “pan” metabolic repertoire, as such finding 49% of the total reactions conserved (**Figure 3a**). The highly conserved metabolic subsystems are aminoacyl-tRNA biosynthesis (100%), lysine biosynthesis (84%), peptidoglycan biosynthesis (78%), glycerophospholipid metabolism (72%) and fatty acid biosynthesis (68%). The pan reactome, on the other hand, was largely represented by carbohydrate and amino acid metabolism, confirming the high diversity of LAB in their ability to catabolize various nutrients as highlighted by comparative genomic analysis. We also investigated the variations in the amino acid auxotrophy (**Figure 3b**) and noticed the absence of several reactions in the relevant biosynthetic pathways (**Figure 3c**).

**Figure 3.**
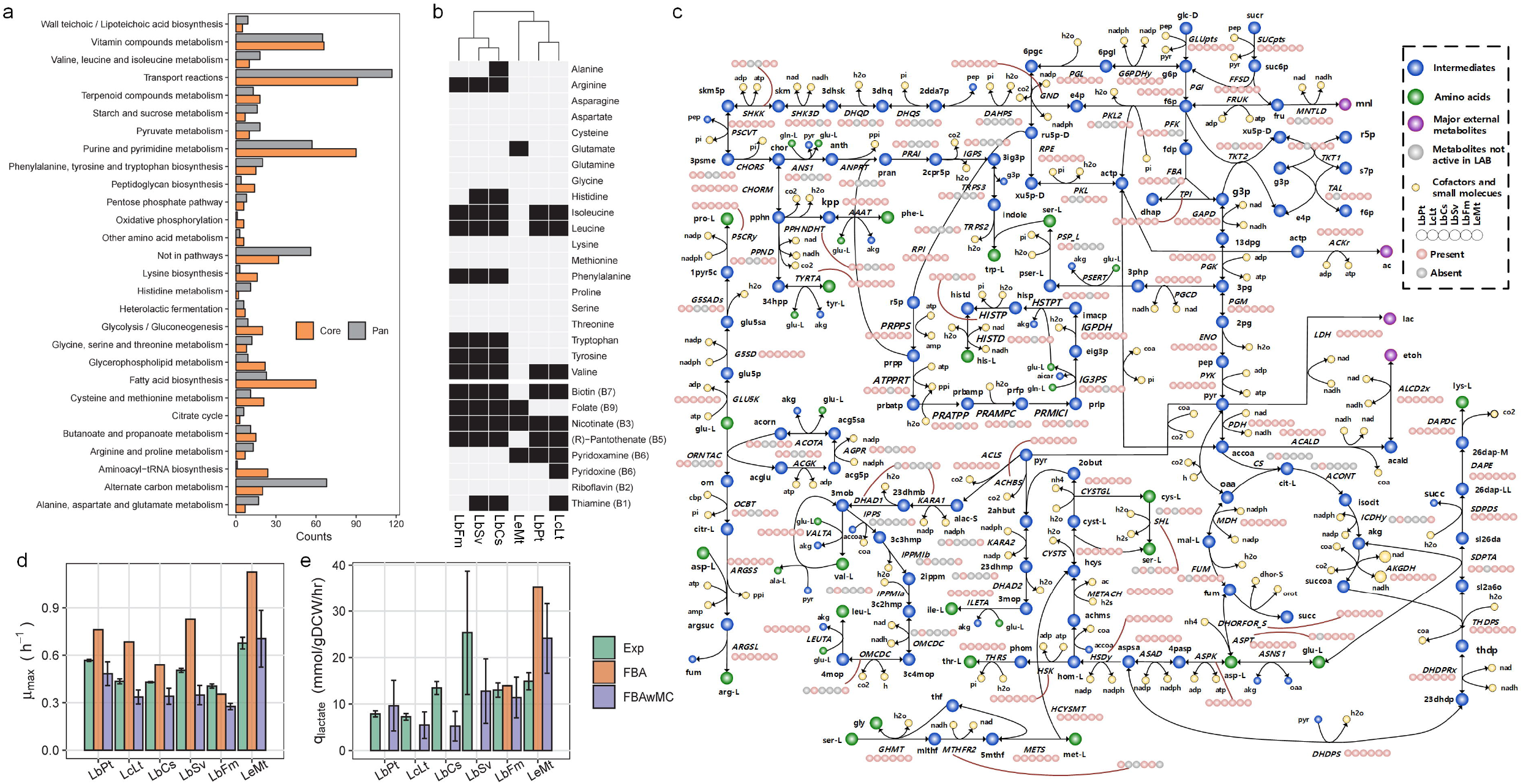
GEM characterized core and pan metabolic capabilities of LAB. (a) Distribution of core and pan reactions in different subsystems, (b) amino acid auxotrophy as revealed by LAB GEMs, (c) metabolic network showing the presence and absence of different reactions in central carbon metabolism and the amino acid biosynthesis pathways, (d) comparison of predicted and measured growth rates, (e) comparison of predicted and measured lactate production rates. The error bars in (d) and (e) denotes the standard deviation of triplicate measurements and the variations in predicted rates over multiple *a_i_* parameters in 5000 samples, respectively.

Following the reconstruction of GEMs, we validated the model predictions using the batch culture data of LAB grown in LABDM (see **Methods**). The initial *in silico* growth rates predicted by flux balance analysis (FBA) were comparable to experimentally measured ones. However, most LAB models failed to simulate the lactate secretion, releasing other fermentative products instead. This motivated us to incorporate additional kinetic constraints for accurately predicting the LAB phenotype. In this regard, implementation of FBA with additional macromolecular crowding constraints (FBAwMC) has been reported to successfully describe the low-yielding acetate overflow in *Escherichia coli*^45^, the Warburg effect in cancer cells^46,47^ and the role of redox cofactors in the switch between low and high yield metabolisms in *Saccharomyces cerevisiae* and LcLt^48^. Therefore, we subsequently formulated an approach based on FBAwMC employing experimentally measured crowding coefficients to investigate the functional metabolic states of each LAB (see **Methods, Supplementary Methods, Supplementary Tables S3-S4 and Supplementary Figures S6-S7**). Remarkably, the growth rates and lactate secretion rates predicted were highly consistent with the culture experiments (**Figures 3d and 3e**), thus confirming that the predominant use of low energy yielding pathways is a hallmark of LAB^42,49^.

### In silico analysis of LAB GEMs highlights the diet-specific patterns in probiotic colonization and postbiotic synthesis

Previously, several cohort-based studies have reported that the gut microbiome composition is more sensitive to the dietary habits than the ethnic or geographical makeup^50–52^. Similarly, we hypothesize that some probiotics may potentially survive and produce beneficial compounds much better in certain diets, which could be tested *in silico*. We explored the probiotic capability of LAB under 11 different diet conditions including vegetarian, vegan, EU average, Mediterranean, DACH, high protein, gluten free, high fibre, diabetes, high fat-low carb, and high fat-high carb by resorting to the model-driven analysis (see **Methods**). The resulting growth patterns signified very distinct diet-specific phenotypes: most of the LAB, especially obligate homofermentative, preferred diabetes and high fibre diets whereas all of them grew very slowly in high fat diets (**Figures 4a-4f** and **Supplementary Figure S8**; p-val < 1 × 10^-150^, Kruskal-Wallis test). The high fibre and diabetes diets are usually rich in inulin type fructooligosaccharides which are catabolized via high ATP yielding pathway involving the (phospho-)fructo-furanosidase (*BfrA* or *SacA*/*ScrB*) enzyme, and thus giving rise to the increased growth rates^53^. LAB do not grow well in high fat-low carb diets since they cannot utilize fatty acids as primary carbon substrates for energy generation although fatty acids are primary constituents for microbial cell walls^54^. Interestingly, we also observed minimal variations in growth rates of heterofermentative LAB (LbFm and LeMt) across different diets, mainly due to the similar levels of ATP produced from diverse carbohydrates via phosphoketolase pathway and their limited capacity to use amino acids as alternative energy sources^37^.

**Figure 4.**
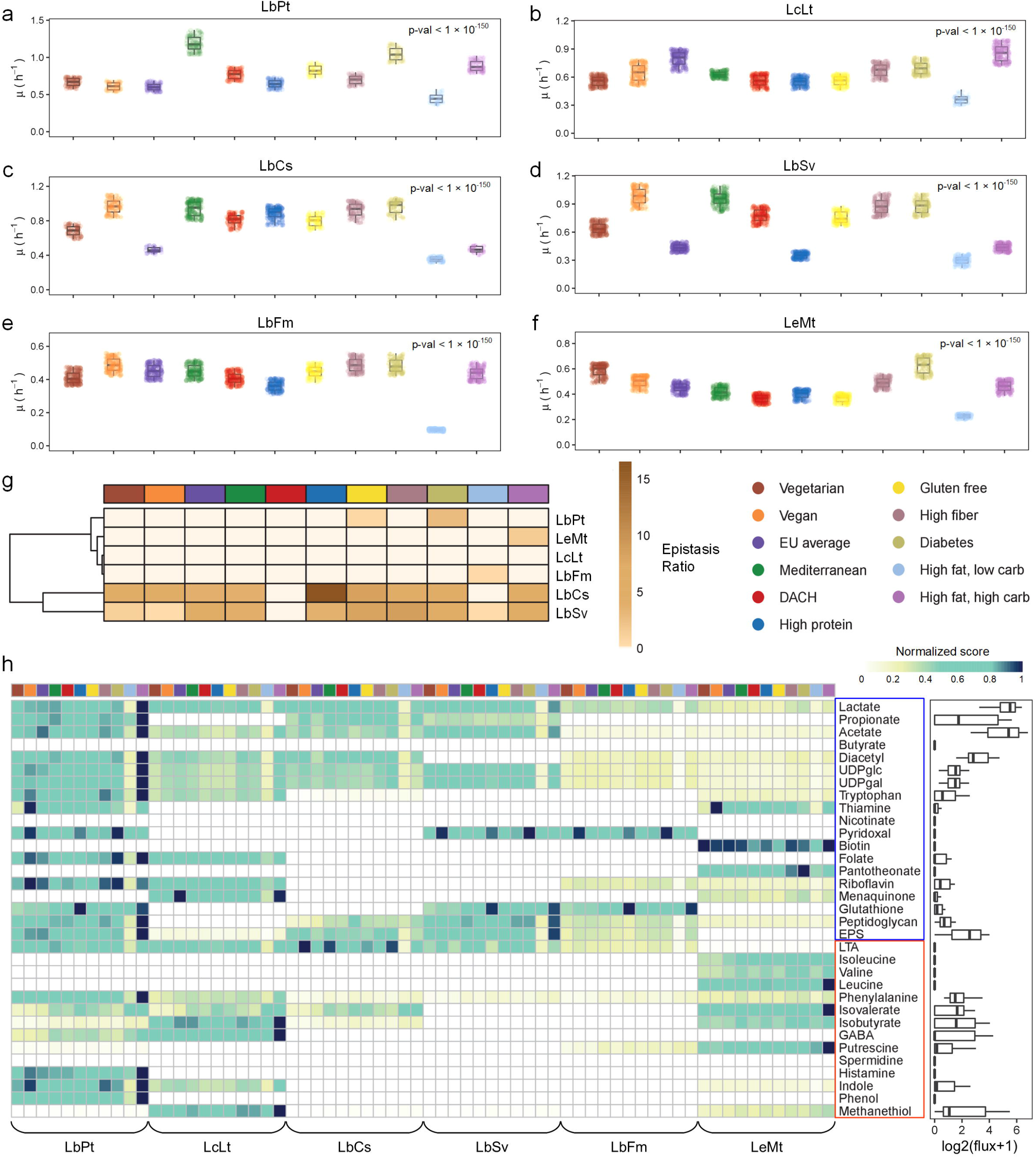
Diet-dependent probiotic characteristics of LAB simulated *in silico*. (a-f) Variations in simulated growth rates across various diets in each LAB species (p-val < 1 × 10^-150^, Kruskal-Wallis test), (g) ratio of positive to negative epistasis and (h) postbiotic production capabilities of LAB in various diets. Growth rates in (a-f) is simulated by constraining each LAB GEM with respective diet conditions over 5000 samples. In (g), the values in heatmap are production potential of each compound normalized to the maximum value across all diets and species. The boxplot in (g) shows the overall variation in production levels across all diets and species. Compounds colored in blue and red denote the beneficial and detrimental postbiotics, respectively.

We subsequently evaluated LAB potentials to produce relevant bioactive compounds (postbiotics) that are beneficial to health, such as the short-chain fatty acids (SCFAs) including butyrate, propionate and acetate. To do so, we computed the theoretical maximum yield of various postbiotic compounds while constraining the biomass flux at 50% of its maximum achievable value (see **Methods** for details). It should be noted that some metabolites secreted by LAB are also known to aggravate health conditions. For example, lipoteichoic acids (LTA) promotes pro-inflammatory responses such as TNF-α induction^61^. Thus, their overall benefit to the host depends on the balance between the production of beneficial and detrimental metabolites. LbPt and LcLt have high amount of growth-coupled production of various beneficial postbiotic compounds such as SCFAs, amino acids and vitamins across all diets comparatively (**Figure 4h**). LbCs and LbSv also secrete similar levels of SCFAs across various diets although they are incapable of synthesizing vitamins and amino acids as high as LbPt and LcLt. On the other hand, LeMt and LbFm synthesized relatively low levels of SCFAs. Similar to the growth patterns, the production level of beneficial metabolites is higher in carbohydrates rich diets, e.g. EU average, DACH, high fibre and diabetes, while it is low in high fat diet. Interestingly, plant-based diets such as vegan, Mediterranean and diabetes consistently enabled several LAB to produce higher amounts of vitamins. Since unique pattern of the gene expression in fermentation pathways was observed for each LAB, we again predicted their postbiotic production with additional constraints based on transcriptome data. These simulations show that while the production of primary metabolites such as lactate and acetate is not significantly affected by gene expression, other compounds are partially regulated at transcript level in few species including LbCs and LeMt (**Supplementary Figure S9**). Here, it should be noted that while we analyzed the biosynthetic capability of LAB, some of them could also consume/degrade the postbiotics including certain amino acids and vitamins synthesized by other LAB as suggested by the auxotrophic analysis (**Figure 3b**), and may partly compromise the beneficial effects.

### Pairwise metabolic interactions of LAB unravel their overall positive associations with commensal species

Probiotics have also shown to selectively interact with the gut flora and alter their abundance^62,63^, particularly in the context of their use as live biotherapeutics. Here, we investigated the pairwise metabolic interactions of each LAB strain with representative probiotics (*Bifidobacterium adolescentis* and other 5 LAB), commensals (*Akkermansia muciniphila, Bacteroides thetaiotaomicron, Enterococcus faecalis, Escherichia coli* W3110, *Faecalibacterium prausnitzii*), opportunistic pathogens (*Pseudomonas putida, H. pylori, Klebsiella pneumoniae, Pseudomonas aeruginosa, Escherichia coli* O157:H7) and pathogens (*Salmonella enterica* ser. typhimurium, *Shigella flexneri, Shigella sonnei, Staphylococcus aureus)* under the 11 diets (see **Methods**). LbCs, LbFm, LbSv and LcLt enhance the growth of commensals while they were either neutral or detrimental to pathogens (**Figure 5, Supplementary Figures S10-S15**). LbCs and LbFm particularly cross-feed *F. prausnitzii* and *A. muciniphilia*, which are widely recognized for their protective role in many metabolic and inflammatory disorders. However, LbPt and LeMt displayed mixed results with no clear distinction between beneficial and detrimental species; they promoted both the growth of commensals such as *F. prausnitzii* and *A. muciniphilia* and pathogenic bacteria such as *Salmonella* and *Shigella*. In addition to strong organism-specific patterns in microbial interactions, we also observed subtle diet-specific patterns. Carbohydrate and protein-rich diets such as Mediterranean, DACH, high protein and high fibre, promoted the growth of commensal and probiotic species in LbCs, LbPt and LeMt (**Figure 5**).

**Figure 5.**
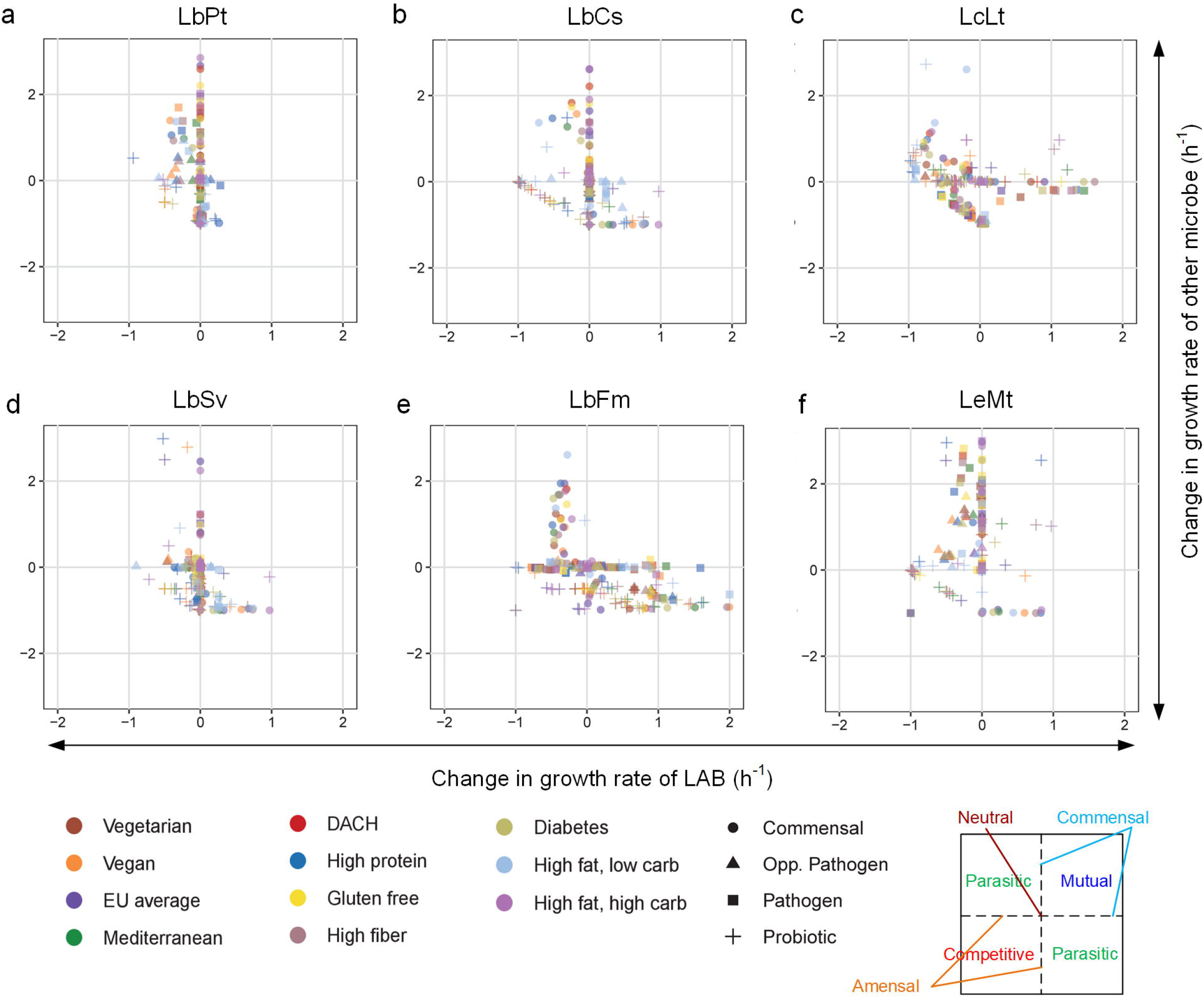
Interactions of LAB with other gut microbes simulated *in silico*. (h-m) outcomes of pairwise metabolic interactions with various other gut microbes. The relative difference in the growth rate of each strain when it grows separately or in combination with another organism is plotted. The interactions of each LAB was evaluated with 5 others and the following representative species: Probiotic – *B. adolescentis*; Commensal – *A. muciniphila, B. thetaiotaomicron, E. faecalis, E. coli* W3110, *F. prausnitzii*; Opp. Pathogen – *P. putida, H. pylori, K. pneumoniae, P. aeruginosa, E. coli* O157:H7; Pathogen – *S. enterica* ser. typhimurium, *S. flexneri, S. sonnei, S. aureus*.

We conducted an extensive literature survey to further validate the simulation results, which showed that LAB could have positive interactions with certain commensal strains. For example, a previous study^64^ reported that the culture supernatants from LcLt and *Lactobacillus paracasei* stimulated *F. prausnitzii* growth by supplementing B-vitamins and lactate/acetate. However, we could not find any evidences of similar interactions between LAB and another keystone gut commensal, *A. muciniphilia*. Therefore, we newly performed cell culture experiments to evaluate such cross-feeding between LAB and *A. muciniphilia, B. thetaiotamicron* and *E. coli* (see **Methods**, **Supplementary Results, Supplementary Figure S16** and **Supplementary Dataset S2**). Among the total 18 combinations experimentally evaluated (6 LAB × 3 commensal strains), 16 of them agreed well with model predictions (F1-score = 0.8333). These results demonstrate the high consistency of model predictions and the utility of GEMs in evaluating microbial metabolite cross-feeding.

## Discussion

While conventional probiotics have been generally developed as food supplements to maintain a good gastrointestinal health, next generation probiotics are expected to provide preventive and therapeutic means to intervene with various diseased states^65^. In this regard, a key challenge is to design combinatorial probiotic formulae based on their synergistic interactions within and with the microbiome. Strain-specific differences and influence of environmental factors including diet could further confound their application^55^. Moreover, there exists no rational framework for screening and functionally classifying probiotics targeting human health, although the Food and Agriculture Organization (FAO) of the United Nations (UN) has currently placed certain guidelines which highly rely on various experimental methods, such as those involving animal models^66^. However, it is impractical to test the efficacy of all candidates and their combinations at the preclinical stage. Therefore, herein we provide a first-pass rational screening procedure to earmark promising candidates using an *in silico* model-guided approach which could be further verified *in vivo*.

Our comparative genomic and transcriptomic analyses suggested LbPt and LbCs as the common LAB with a higher proportion of functionally relevant genes supporting gastrointestinal survival via the resistance to acid, bile and oxidative stress, production of bacteriocins and CEPs, and mucosal adhesion. Interestingly, both LAB have relatively larger genomes and coincidentally belong to the facultative heterofermentative group. Another interesting finding is the extraordinary diversity of LAB pan genome, especially enriched on carbohydrate and amino acid metabolism, which motivated us to explore the functional metabolic capabilities as well as the cellular fitness across different diet conditions. Indeed, our model-driven analysis confirmed diet-specific patterns where high fat-low carb (ketogenic) diets lead to poor probiotic performance and diets rich in inulin type fructooligosaccharides is preferable, in general. Consistent with our results, recent clinical studies have also shown that diets rich in high fat-low carb cause unfavourable changes in gut microbiome and reduction in fecal SCFAs which could be due to the elimination of probiotic bacteria^67–69^. These observations clearly demonstrate the need to either select appropriate probiotics depending on the dietary habit of a person or administer them as synbiotics, i.e., probiotics with relevant ‘prebiotics’, to enable their efficient functioning in gut. Such rational administration of probiotics becomes even more significant for live biotherapeutics. In this regard, although our results indicate LbPt, LcLt and LeMt to have highest capabilities for the postbiotic production, they also synthesize increased levels of LTA which is highly implicated in the stimulation of pro-inflammatory cytokines^61^, and thus caution must be exercised while recommending them to patients with inflammatory bowel disease (IBD). Our simulations also hinted that LbCs and LbFm could be promising probiotic candidates to treat/manage IBD as they both selectively promoted the growth of two commensals (*A. muciniphilia* and *F. prausnitzii)* that are reported to have a strong inverse correlation with IBD^70^. Numerous earlier studies have reported that LbCs strains consistently confer anti-inflammatory and protective effects against IBD and colorectal cancer^71–76^. On the other hand, LbPt and LeMt displayed positive interactions with both pathogens and commensals growth, and thus may preferably be avoided for treating immunocompromised patients whose natural ability to eliminate gastrointestinal pathogens is generally limited^77^. These observations demonstrate the advantage of adopting a multi-omics based systems approach compared to the widely used comparative genomics analysis.

In summary, the current study highlights that the fitness and postbiotic synthesis capabilities of probiotics are diet-dependent while their interactions with gut microbes are largely organism-specific. Our *in silico* simulation results clearly indicate that while high fiber diets promote the probiotic growth and functionality, high fat diets lead to poor performance, which is in high concordance with previously published clinical data^67–69^. However, we urge caution in extrapolating inferences from the presented framework as the annotation of microbial genomes vary considerably. In addition, some of the probiotic evaluation aspects, e.g. colonization, are still uncertain^55^, which can be simply addressed by further revising the presented *in silico* analysis. Despite such restraints, the systematic framework proposed here plugs the gap between comparative genomic analyses and *in vivo* evaluation. It also seeds the foundation for model-guided selection of relevant probiotics, and can be further extended by accounting relevant confounding factors in clinical settings, such as host-genetics and actual diets, through community-wide efforts to foresee well-controlled clinical trials. Overall, the proposed approach is poised to accelerate the characterization of probiotic capabilities of multiple candidates beyond the conventional *Lactobacillus* and *Bifidobacterium* species, thus speeding up the rational design of smart probiotics in near future.

## Methods

### LAB strains, media and growth conditions

LAB strains, *Leuconostoc mesenteroides* subsp. mesenteroides ATCC 8293, *Lactobacillus casei* subsp. casei ATCC 393, *Lactobacillus plantarum* WCSF1, *Lactobacillus salivarius* ATCC 11741, *Lactobacillus fermentum* ATCC 14931 were from ATCC, and *Lactococcus lactis* subsp. cremoris NZ9000 was from Boca Scientific. The lyophilized cultures were first revived by aseptically transferring the contents of the ampoule into 10ml each of MRS and GM17 media, equally. The revived cultures were then sub-cultured using the same media to ensure stability. A chemically defined medium capable of supporting all six LAB strains (LABDM) was developed by iteratively modifying components of a previously established CDM^36^ (see **Supplementary Dataset S1**). Seed LAB cultures for fermentation experiments were prepared by transferring the 10% (v/v) cultures to LABDM and incubating them overnight at 30°C. Biological triplicates of the LAB were grown in 50 mL conical tissue culture tubes (Thermo Fisher Scientific) as static cultures with added reducing agent, sodium thioglycolate (1g/L), and no head-space to maintain anaerobic conditions. A separate culture tube was maintained for each time point, to avoid aeration during sampling.

### Growth and biochemical analysis of culture supernatants

Cells and supernatants were harvested from the biological triplicate LAB cultures cultivated in LABDM at regular time intervals spanning the exponential growth phase for various growth and other biochemical assays. Optical density at 600nm (OD600) was measured using Shimadzu UV-1700 spectrophotometer. Dry cell weights (DCW) were obtained by centrifuging known volume of cell cultures, subsequently drying the pellet at 100°C overnight, and weighing them using a balance. Standard curves of OD600 vs gDCW were plotted to estimate the conversion factors. Glucose and lactate profiles at different time intervals spanning exponential phase were analyzed using YSI® biochemistry analyzer. Amino acid profiles were measured using Waters ACQUITY-UPLC system, AccQ·Tag™ Ultra Column (2.1 × 100 mm) and AccQ-Tag derivatization kit, following the manufacturer instructions. Note that biological triplicates were employed for all sample measurements.

### Cell volume and cell number measurements

LAB single cell volumes were estimated by approximating the shapes of LcLt and LeMt to spherical, and LbPt, LbCs, LbSv and LbFm to cylindrical dimensions. Average cell diameters and lengths of at least a hundred cells measured using an optical microscope at 100X magnification under oil immersion were used to estimate single cell volumes. Bacterial cultures were appropriately diluted before microscopic measurements to ensure proper dispersal of cells. Cell numbers in a given volume of the same diluted culture were enumerated using plating and automatic CFU counter (Scan 1200, Interscience, Saint Nom, France).

### Determination of total protein content

Total protein content was measured using Bradford assay^81^. Briefly, cells pellet obtained from culture of known volume and OD600 were suspended in 1 mL ice cold lysis buffer containing 10 mM Tris-HCL, pH 8.0, 1 mM EDTA, and 100 ul of lysozyme (10 mg/mL), and incubated on ice for 15 min. Next, the cells were sonicated at a frequency of 20 kHz for 10 × 30 sec with 1 min interval between each sonication cycle. Ten rounds of sonication resulted in “close to complete” protein release, after which the protein concentration saturated. Bradford protein assay kit (Bio-Rad®) with standard Bovine Serum Albumin (BSA) solution was used to estimate the protein concentration in the sonicated samples after sufficient dilution, which were then converted to total protein content per gram DCW using appropriate conversion and dilution factors.

### Total RNA extraction and purification

To extract the total RNA, 10 mL aliquots were harvested from the biological triplicate LAB cultures cultivated in LABDM at time points that correspond to their respective mid-exponential phases. It was first mixed with 10 mL of Qiagen RNAprotect reagent and incubated for 10 min at ambient temperature to stabilize RNA. Bench and biosafety cabinet surfaces used for RNA work were decontaminated with RNaseZAP (Thermo Fisher Scientific) to minimize contact of samples with RNase. Half of the sample aliquotes (10 mL) containing RNAprotect reagent were centrifuged at 6000 x g for 10 min. The remaining samples were stored at −80 °C up to 4 weeks for any later use. Prior to RNA extraction, the cell pellets were treated with 1 mL icecold lysis buffer containing 10 mM Tris-HCL, pH 8.0, 1 mM EDTA, and 100 uL of lysozyme (10 mg/mL), all prepared using RNase-free water, and incubated for 30 min. The lysed solution was further subjected to mechanical cell disruption involving nuclease-free glass beads and thermomixer to ensure maximum release of RNA. Qiagen RNeasy Mini kit was used to extract total RNA from the cell pellets following manufacturer instructions. DNase treatment was performed using Turbo DNA-free kit (Thermo Fisher Scientific) to remove any genomic DNA contamination. Integrity and quality of the extracted RNA was initially checked by the presence of 16s and 23s rRNA in samples subjected to agarose gel electrophoresis and later confirmed using Agilent Bioanalyzer. Bacterial Ribo-Zero Magnetic kit (Illumina) was used to deplete rRNA and enrich mRNA in the samples, following manufacturer’s instructions. Agilent Bioanalyzer was used to ensure the depletion of rRNA from all samples before the preparation of cDNA libraries.

### Preparation of cDNA libraries and RNA-sequencing

Fragmentation of rRNA depleted mRNA samples and subsequent preparation of cDNA libraries were performed using Illumina TruSeq Stranded mRNA Library Preparation kit (Low Sample Protocol), following manufacturer’s instructions. First cDNA strand synthesis in the reverse transcription polymerase chain reaction (RT-PCR) was performed using the reagents supplied by the manufacturer along with the SuperScript II reverse transcriptase (Thermo Fisher Scientific). Indexed cDNA libraries were pooled and single-end sequenced on Illumina HiSeq 2500 Rapid V2 platform to 51 bp read length.

### Commensal culture with LAB supernatants

Brain heart infusion (BHI) broth supplemented with heme, N-acetylglucosamine was used to culture all three commensal strains (*A. muciniphila* ATCC BAA-835, *B. thetaiotaomicron* VPI-5482, and *E. coli* W3110). Media were purged with N2 gas prior to use, ensuring they are devoid of oxygen. LAB strains were first statically grown in CDM overnight at 30°C. Simultaneously, commensal bacteria were inoculated to BHI medium and allowed them to grow overnight at 37°C. Supernatants from the LAB culture 10% v/v was then transferred to freshly prepared BHI in a culture tube. The overnight-grown commensal culture was seeded to this tube to a final OD600 of 0.1-0.2. All experiments were conducted in duplicates and were done in an anaerobic chamber. The culture tube was placed inside anaerobic jar and purged with N2 for 10 minutes before including the GazPak EZ sachets (Becton-Dickinson). After sealing, the jar was placed in a 37°C incubator. Growth of the commensal bacteria was recorded by measuring OD after 12 hours of incubation.

### Statistical analysis of LAB growth characteristics

LAB growth characteristics were evaluated on the basis of three parameters: maximum growth rate, time taken to reach mid-exponential phase and the biomass yield over glucose. Briefly, maximum growth rate is the maximum value of slope along the log transformed growth curve. Time to mid-exponential is the time required to reach half the maximum OD in the culture. Biomass yield over glucose is the ratio of LAB gDCW accumulated over the culture to that of g glucose consumed. Maximum growth rate and time to mid-exponential were calculated by fitting a logistic equation to the culture data using Growthcurver^82^.

### Comparative genomic analysis

#### Reconstruction of phylogenetic tree

To understand the evolutionary perspective of LAB strains examined in this study, phylogenetic tree was constructed from the whole genome sequences using Type (Strain) Genome Server (TYGS)^83^. Note that we also included four other LAB strains, *Lactobacillus reuteri* ATCC 53608, *Lactobacillus delbrueckii* subsp. *bulgaricus* ATCC 11842, *Lactobacillus acidophilus* NCFM, and *Bacillus subtilis* subsp. *subtilis* str. 168, for this particular analysis. Whole genome phylogeny trees were reconstructed based on Genome BLAST Distance Phylogeny (GBDP) approach where all genome sequences were compared pairwise and intergenomic distances was calculated using the ‘trimming’ algorithm and distance formula d5^84^. Phylogenetic trees with branch support were then inferred from intergenomic distances using FastME^85^ based on the original and pseudo-bootstrapped matrices, and visualized using Interactive Tree Of Life (iTOL)^86^.

#### Identification and functional annotation of LAB “core” and “pan” genome

In order to identify the “core” and “pan” genome, we first established the homology relationships between any pair of LAB species using InParanoid^87^. Since InParanoid orthology detection is based on pairwise comparisons, we generated 15 InParanoid tables, identifying the orthologous genes for each genome pairs. Subsequently, we compared the list of one-to-one orthologs across the LAB using LbPt as reference, and established the “core” genes that are present in all species. The remaining non-unique gene count in each LAB is considered as “pan” genome. Functional enrichment of “core” and “pan” genome was performed using eggNOG mapper based on the NOG categories^88^. Note that NOG categories are similar to well-known COG categories but it is identified in an unsupervised manner.

#### Genomic comparisons for non-metabolic probiotic determinants

BLASTp orthology search was used to mine the LAB genomes for identifying ORFs encoding proteins for resisting acid, oxidative and bile stresses, protein lysing proteases and anchor cells on host mucosal layer surfaces. To do so, we first surveyed for genes known to be associated with probiotic characteristics in various LAB (See Supplementary Information). Appropriate cutoffs (Identity > 50%, query coverage > 70%, E-value < 1E-06)^89^ were used to select orthologs. Potential bacteriocin operons in the LAB genome was mined using BAGEL^90^.

### Transcriptome data analysis

#### Transcriptome assembly and gene expression quantification

First, the quality of RNA-seq reads were assessed with FastQC^91^. The adapters and low-quality reads were subsequently trimmed using Trimmomatic v0.32^92^ and the trimmed reads were then aligned with the respective genome assemblies using STAR v 2.5.3a^93^. Finally, RSEM v1.3.0^94^ was used to quantify expression levels in terms of counts, FPKM and TPM from the alignment files. ***Functional enrichment of gene expression categories*:** First, the genes are grouped into three groups based on TPM values: i) low expression range (log2(TPM+1) < 2.5), ii) medium expression range (log2(TPM+1) > 2.5 & < 10) and iii) high expression range (log2(TPM+1) > 10). Subsequently, a p-value is calculated representing the significance of NOG category enrichment in different gene expression groups using the Fisher’s exact test. The p-values was subsequently adjusted for false-discovery rates based on previously proposed method^95^. ***Comparison of gene expression in orthologous genes*:** The direct comparison of gene expression across LAB species in gene wise manner is not possible because their genome sizes vary as well as their genome functionalities are different. Therefore, in order to compare the gene expression levels of same gene functions across LAB, we first identified one-to-one orthologues as mentioned previously. We subsequently normalized the raw counts of these orthologues based on a set of housekeeping genes using RUVseq^96^. The housekeeping genes were identified using the following criteria: i) genes which have expression over 95 percentile range in each LAB and ii) genes which are present in one-to-one orthologues list. Orthologous gene expression profiles were then compared using PCA and Spearman correlation in a pairwise manner.

#### Identification of differentially expressed genes

DEseq2^97^ was used to identify differentially expressed one-to-one orthologous genes across the 6 LAB. Note that we used the gene lengths in each LAB as an additional normalization factor in this step since the gene length will be different in each LAB for the same gene. Those genes with FDR adjusted p-values < 0.01 were classified as significantly differentially expressed genes.

#### Functional enrichment of differentially expressed genes

Differentially expressed genes were functionally enriched based on KEGG pathways and GO terms using Database for Visualization and Integrative Discovery (DAVID)^98^. Pathways and GO terms which have an FDR adjusted p-value < 0.01 were only considered for further analysis.

### Reconstruction of LAB genome-scale models

GEMs of LbSv, LbFm and LbCs were newly developed based on their respective genome annotations, whereas LbPt, LeMt and LcLt GEMs have been updated from the previously published ones^37,42,43^. The reconstruction/updating of GEMs involved standard procedures, which can be found elsewhere^44^.

To newly reconstruct GEMs, initially the metabolic pathway information based on genome annotations of the LAB, was collected from databases, including KEGG^99^ and MetaCyc^100^. The draft reaction networks were then assembled by combining information from these databases, along with the information based on gene orthologs of the existing LAB models^37,42,43^. Next, an artificial reaction, known as biomass reaction, which is typically used as an objective function in constraint-based flux analysis to predict metabolic fluxes of exponential growing cells^101^, was added to each newly developed model based on experimental and literature information (see **Supplementary Dataset S3**). Note that since protein constitutes the major macromolecular component of biomass, we measured the total protein content of different LAB experimentally and used it for biomass reaction formulations (**Supplementary Dataset S3**). Gaps in the these draft networks were then identified using gapFind algorithm^102^ and subsequently filled by adding new reactions based on bibliographic information. Subsequently, we formulated the gene-protein-reaction (GPR) relationships and rectified the model inconsistencies, including elemental, charge imbalances. We also used the amino acid auxotrophy data available in literature to manually curate the reconstructions by reducing the inconsistencies in auxotrophies observed in experiments and during constraint-based simulations. The fermentable substrate phenotyping data for 14 different carbohydrates was then used to test the agreement between model predictions and experiments, subsequently eliminating the inconsistencies through the addition of relevant metabolic and transport reactions. Growth and non-growth associated ATP maintenance costs were either adopted from existing LAB genome-scale metabolic models or estimated based on available culture data (**Supplementary Dataset S3**). Finally, the reconstructed models’ quality and consistency were evaluated using online tool MEMOTE^103^ (**Supplementary Dataset S4**)

Existing GEMs were updated by first identifying the network gaps using gapFind algorithm^102^ and their subsequent resolution by adding new reactions based on bibliographic information. Next, we included new genes, reactions and metabolites and updated the GPR for several existing reactions based on the latest genome annotations, following the standard procedures^44^. Finally, similar to the new GEMs, we also used the amino acid auxotrophy data available in literature to manually curate the reconstructions and evaluated the model quality and consistency using MEMOTE.

### Constraint-based flux analysis with macromolecular crowding constraints

Constraint-based flux analysis with macromolecular crowding constraints^45^ was used to analyze the metabolic phenotype of the LAB under various environmental conditions such as

LABDM and different dietary regimes. The corresponding optimization problem can be mathematically represented as follows:

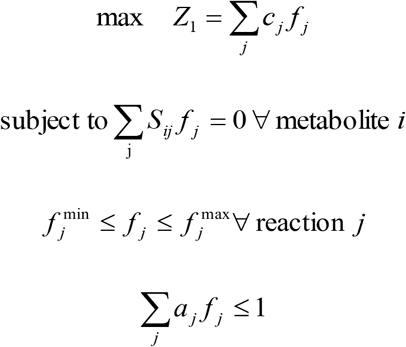

where *Z* is the cellular objective, *c_j_* is the relative weights of each metabolic reaction to biomass formation. *S_ij_* is the stoichiometric coefficient of metabolite *i* of reaction *j*; *f_j_* is the flux through the reaction *j*; 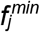 and 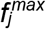 are lower and upper bounds on the flux through the reaction *j*, respectively; and *a_j_* is the ‘crowding coefficient’ which need to be calculated based enzyme kinetic parameters, i.e. turnover numbers, and cytoplasmic density (see **Supplementary Methods**).

In order to simulate the physiology of exponentially growing LAB in LABDM, the biomass reaction was maximized while simultaneously constraining the uptake/secretion rates of glucose and all 20 amino acids at experimentally measured values. Similarly, the exchange reactions were constrained at corresponding uptake rates provided in VMH database^104^ for simulating the LAB growth phenotype under various diet conditions. Note that we used 5000 different permutations in each simulation to account for reactions with unknown C values and obtained 5000 flux solutions. The geometric mean of the flux solutions surrounding the maxima of the resultant lognormal distributions accurately represent the average cellular state^105^, and were then used to assess the phenotype (growth rate/byproduct secretion) of the LAB (see **Supplementary Results** and **Supplementary Figure S5**). All simulations were implemented using COBRA toolbox^106^ and Gurobi7 (http://www.gurobi.com) optimization solver (see **Supplementary Methods** for more details).

### Simulation of postbiotic production potential in LAB

In order to compute the theoretical yield of the postbiotic metabolites, pseudo-reactions representing their secretion was added and iteratively used as the objective function. Biomass was constrained to 50% of the optimum value. FBAwMC was implemented using COBRA toolbox^108^ functionalities and the objective was maximized to obtain the *in silico* postbiotic yields of each LAB while constraining the exchange reactions at corresponding uptake rates provided in VMH database for each diet. We used 5000 different permutations for each FBAwMC simulation to account for reactions with unknown C values. The geometric mean of the resulting 5000 flux solutions surrounding the maxima of the lognormal distributions accurately represents the average cellular state^105^, and was then used to calculate the postbiotic yield. Finally, the yield of each postbiotic compound was normalized to the maximum value across different diets and LAB.

### In silico examination of pairwise interactions between LAB and other microbes

Pairwise microbial interactions between LAB and any other microbe under anaerobic conditions were evaluated *in silico* based a method proposed earlier^109^. Briefly, the LAB model was first paired with any of other microbe GEMs^110-12^> by introducing a common lumen compartment to facilitate the exchange of nutrients between them. Second, the exchange reactions of various compounds in each diets were added to the lumen.

The pairwise interactions were then evaluated using the enzyme-capacity constrained paired model as follows. First, the biomass of both species in the paired model was maximized under a particular diet or BHI media to obtain the growth rates of individual species when they grow together. Next, the maximum growth of individual species when they grow in isolation under the same diet was computed while simultaneously constraining all the reactions of other species to zero. Finally, by comparing the growth rates of paired and isolation conditions, the pairwise interactions are classified as follows: i) competition – if both organisms grow slower in paired condition than isolated conditions, ii) parasitism – one species grows faster while other grows slower in paired condition than isolated conditions, iii) mutualism – both species grow faster in paired condition than isolated conditions, iv) amensalism – one species grow slower while the other is unaffected in paired condition than isolated conditions, v) commensalism – one species grow faster while the other is unaffected in paired condition than isolated conditions and vi) neutralism – no appreciable difference in both species growths between paired and isolated conditions. Note that a species was considered to grow faster in the paired condition when its growth rate was more than 10% higher than isolated condition. The exchange reactions were constrained at corresponding uptake rates provided in VMH database^104^ or values calculated based on a previous study^121^ for simulating the LAB growth phenotype under various diet conditions or BHI media, respectively. All pairwise interactions were simulated using microbiome modeling toolbox^122^.

## Acknowledgements

The authors thank Shawn Hoon and Fong Tian Wong for advice on RNA-seq library preparation steps, Elena Heng for experimental assistance in transcriptome sequencing, and Say Kong Ng and Tessa Tan for their advice and support on HPLC analysis.

## Author contributions

L.K., M.L., D.S.-W.O. and D.-Y.L. conceived the project. L.K., P.-Y.L. and M.B. performed LAB experiments, and profiled culture supernatants. P. L. H. and D.-S. P performed commensal culture experiments. L.K. formulated the new LABDM with inputs from D.S.-W.O. L.K., P.-Y.L. and M.L. were involved in transcriptome sequencing, assembly, and relevant bioinformatics analysis. L.K. and M.L. performed the comparative genomic analyses. L.K. and M.L. reconstructed the genome-scale models, developed in silico methods and implemented them. L.K. and M.L. drafted the initial manuscript. L.K., M.L., D.S.-W.O. and D.-Y.L. were involved in editing and revising the manuscript. D.S.-W.O. and D.-Y.L. supervised and coordinated the project.

---

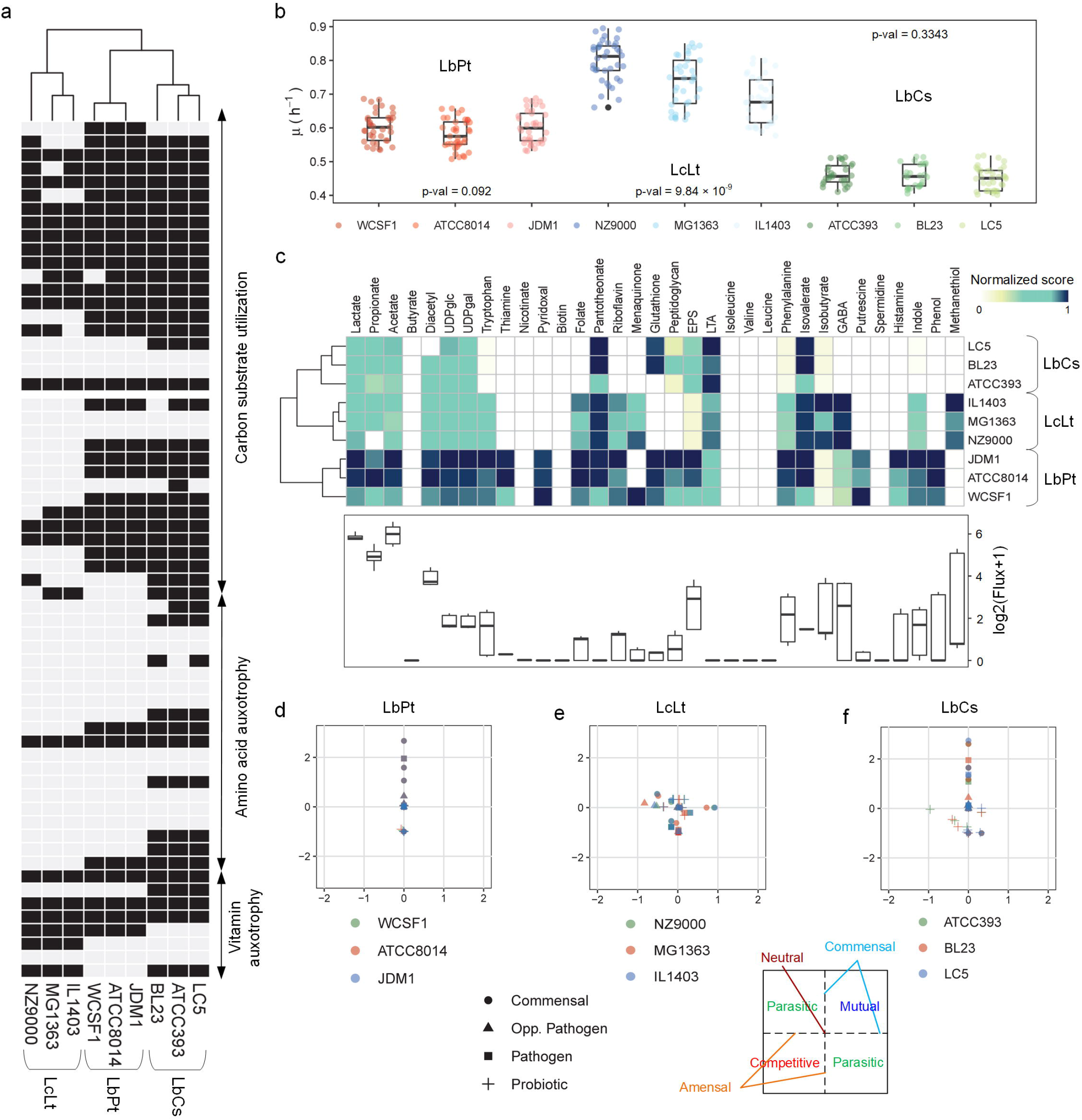

