## Supplementary Text for "Systematic evaluation of genome-wide metabolic landscapes in lactic acid bacteria reveals diet-induced and strain-specific probiotic idiosyncrasies"

### Supplementary Results

#### *Comparative genomics analysis of non-metabolic probiotic determinants*

Apart from the metabolic flexibility, LAB also synthesize a variety of proteins for resisting acid, oxidative and bile stresses, conferring anti-microbial activities, protein lysing proteases and to anchor cells on host mucosal layer surfaces. Since all these proteins are critical for a probiotic organism to colonize and sustain the harsh environments of human gut, we mined the genomes of all 6 LAB species to identify relevant proteins (see **Methods**). Notably, all LAB encode for multiple cell adhesion factors which could help them attach in the mucosal layers. In stress resistance category, while all LAB encode at least one protein, some of them have more than one (**Figure 1g**). For example, LbPt genome contain five different bile resistance genes and four unique acid resistance genes. Many LAB also produce and secrete anti-microbial peptides, known as bacteriocins, which can be broadly classified into three classes: i) class I bacteriocins – small heat stable posttranslationally modified peptides, ii) class II bacteriocins – small heat stable posttranslationally unmodified peptides and iii) class III bacteriocins – slightly large tthermo-labile stable peptides[1]. We identified 8 genes encoding for potential class II bacteriocins in LbPt, and one each in LeMt and LbCs (**Figure 1g**). We also identified a potential gene encoding for the class III bacteriocin in LbSv. The ability to release cell envelope proteases (CEPs) which cleave the casein peptides in milk environment into shorter peptides/amino acids for utilizing them as nutrient sources is also a primary attribute of most LAB. Additionally, some LAB CEPs are shown to alter the inflammatory bowel disease by acting upon the inflammatory mediators. So, we also mined all the LAB genomes and identified that all of them could potentially produce at least one such CEP (**Figure 1g**).

#### *LAB GEM consistency evaluation using MEMOTE*

All nine GEMs reconstructed in this study were also evaluated using MEMOTE[2] for their consistency and completeness. The MEMOTE reports of all LAB models are provided in

**Supplementary Datset S4**. While the overall MEMOTE scores are around 50%, all 9 GEMs had a very high consistency score (> 90%), indicating that the models have good stoichiometry accuracy, proper mass balances and charge balances, and do not have high number of reaction cycles. The low overall scores were mainly because of the missing annotations which may slightly affect their use in automated tools but not compromised in model quality.

***FBAwMC predict bimodal lognormally distributed growth rates in LAB***

Utility of LAB GEMs to evaluate the quantitative aspects of the metabolism relies on their predictive power. The production of lactate has been observed as the most important characteristic in LAB, having prominent role in their evolution[3]. Moreover, lactate biosynthesis is tightly linked to the energy and redox metabolism of LAB, suggesting the need for accurately predicting lactate production in order to better capture their metabolic states. However, when simulated with FBA, the LAB models fail to predict low yield metabolism characterized by lactic acid production[4,5], suggesting the measurement of lactic acid an essential step to derive reasonable *in silico* predictions. In this regard, implementation of FBA with additional constraints based on macromolecular crowding (FBAwMC) has shown to successfully predict the low-yielding acetate overflow in *E. coli*[6] and the Warburg effect in cancer cells[7,8]. The method has later been applied to *Saccharomyces cerevisiae* and LcLt to explain the role of redox cofactors such as NADH in the switch between low and high yield metabolism[9]. Therefore, here we used FBAwMC (see **Supplementary Methods**) to predict the low yield metabolism in all six LAB.

The implementation required measurement estimation of crowding coefficients for each enzyme/reaction, which in turn needed experimental measurement of parameters such as the average cell sizes and their correlation to dry weight for each LAB (see **Methods**). The estimation of crowding coefficients requires enzyme kinetic information, especially turnover numbers, and the availability of such data is limited only to a subset of well-studied enzymes that are part of GEMs. Hence, the crowding coefficients of reactions for which no data was available were randomly assigned from the list of coefficients for enzymes whose kinetic data was available. The typical growth rate predictions obtained for ~5000 samples generated for each LAB based on such random assignments are shown in **Supplementary Figure S5**. Interestingly, the frequency of growth rates in different samples followed bimodal lognormal distribution for all LAB. Considering growth as the primary objective of a cell and hence correlation between the overall gene expression levels and growth, the gene expression levels across the samples are also expected to follow a lognormal distribution. Remarkably, similar conclusions were made from experimental studies involving single-cell transcriptomics and flow cytometry, i.e. the gene expression levels in cell population growing together follow log-normal distribution[10–12]. Hence, the two geometric-mean growth rates ideally represent their most probable low- and high-yield metabolic states for the given environmental condition.

***Differences in the metabolic and postbiotic capabilities of LAB strains***

A vast amount of literature has previously reported the considerable differences in probiotic capabilities across various strains in the same species[13–16]. Therefore, here, we evaluated the metabolic and probiotic capabilities of 3 strains of 3 LAB species, LbPt, LcLt and LbCs, to assess whether such differences are as high as the ones observed between the different species. For this purpose, we first reconstructed the corresponding strain-specific GEMs from the representative strains using orthologous genes. We further curated each strain-specific model by accounting for its unique genome content (see **Methods** for detailed reconstruction procedure). Following the reconstruction of strain-specific GEMs, we first compared the metabolic capabilities of different strains by examining their ability to ferment 32 different carbon substrates, and auxotrophy requirements to 20 amino acids and 8 vitamins. These results clearly indicated that while there exist some differences across the strains, they were very similar to other strains in same species than that of other species (**Figure 6a**).

We next compared how the different strains perform in the "EU average" diet in terms of cellular fitness and metabolic interactions with other representative species in human gut. Overall, the maximum achievable growth rates of different strains of same species were very similar (**Figure 6b**), possibly due to the use of same biomass coefficients across strains due to lack of strain-specific macromolecular composition data availability. We also compared the potential strain-specific differences in terms of postbiotic producing abilities under EU average diet. These results also indicate while the strain specific differences in postbiotic synthesis are minimal when compared across species (**Figure 6f**). However, the differences in LbCs strains where much more pronounced when compared to LbPt and LcLt. Particularly, the yields of propionate, diacetyl, UDP-glucose, UDP-galactose, LTA and peptidoglycan were markedly different across LbCs strains.

Finally, we analysed the pairwise interactions of different LAB strains with representative gut microbes in the "EU average" diet to evaluate whether different strains elicit dissimilar interactions or not. Unlike cellular growth, the pairwise interactions indeed showed notable differences across strains of same LAB, particularly in LbCs (**Figure 6c-e**). For example, the ATCC393 and BL23 strains of LbCs are mutually associated with *B. thetaiotaomicron*, whereas the LC5 strain display a commensal relationship. Similarly, the ATCC393 and BL23 strains are parasitic to several pathogens whereas LC5 show amensalism interaction with them.

***Spent-medium transfer experiments to validate in silico pairwise interactions***

In order to test the validity of the *in silico* pairwise interaction outcomes, we performed spent-medium transfer experiments for all 6 LAB and 3 commensals, *A. muciniphilia*, *B. thetaiotaomicron* and *E. coli*, grown in Brain-heart infusion (BHI) medium (see **Methods**). The growth of the three commensals with and without the cell-free supernatant from each LAB was evaluated (**Supplementary Figure S15A-C**). We also simulated the pairwise interactions between each LAB and the three commensals in BHI media under anaerobic conditions (see

**Methods**). The fold change of commensal growth with and without LAB supernatants was then calculated and compared with the corresponding *in silico* predicted fold-change change in commensal's growth between pairwise and single cultures (**see Supplementary Dataset S3**). Although the growth changes were minor, we observed the direction of change (growth-promoting or growth-inhibiting) to be consistent with *in silico* results (Overall accuracy=89%). In particular, model predictions were highly consistent with experimentally observed growth promoting and inhibiting trends of most LAB with *A. muciniphilia* and *B. thetaiotaomicron*, respectively. The effect of cell-free supernatants of *L. casei* and *L. plantarum* on *E. coli* growth showed trends opposite to corresponding model predictions. Such, inconsistencies could be attributed to unknown exchanges between the LAB and commensals that are not accounted in models, hinting at the scope for further curation of the LAB and commensal GEMs. Moreover, cross-feeding may only partly capture the microbial interactions, where other interactions such as physical cell-cell contact and signalling mechanism could also play a role in deciding the fate of such interactions. However, the spent-medium transfer design has captured the qualitative aspects predicted by the models underlines the importance of nutrient exchanges in governing LAB-commensal interactions.

### Supplementary Methods

#### *Constraint-based flux analysis with macromolecular crowding constraints*

Constraint-based flux analysis was used to analyze the metabolic phenotype of the LAB under various environmental conditions such as LABDM and different dietary regimes. The corresponding optimization problem can be mathematically represented as follows:

$$\max Z = \sum_j c_j f_j \qquad (1)$$

$$\text{subject to } \sum_j S_{ij} f_j = 0 \ \forall \text{ metabolite } i \qquad (2)$$

$$f_j^{min} \leq f_j \leq f_j^{max} \ \forall \text{ reaction } j \qquad (3)$$

where, $Z$ is the cellular objective, $c_j$ is the relative weights of each metabolic reaction to biomass formation. $S_{ij}$ is the stoichiometric coefficient of metabolite $i$ of reaction $j$; $f_j$ is the flux through the reaction $j$; $f_j^{min}$ and $f_j^{max}$ are lower and upper bounds on the flux through the reaction $j$, respectively.

Additional constraints based on macromolecular crowding is

$$\sum_{j=1}^{N} v_j n_j \leq V \qquad (4)$$

Molecular crowding constraint, $v_i$ is the volume of macromolecule (enzyme '$j$'); $n_j$ is the number of moles of the $j^{th}$ enzyme; $V$ is the volume of the cell. Dividing both sides by cell mass (M) and taking into account the enzyme concentration, $E_j = n_j/M$ (moles/unit Mass), we obtain the constraint on maximum attainable enzyme concentrations.

$$\sum_{j=1}^{N} v_j E_j \leq \frac{V}{M} \qquad (5)$$

$$\sum_{j=1}^{N} v_j E_j \leq \frac{1}{C} \qquad (6)$$

where, $C = M/V$, is the cytoplasmic density.

Since the maximum amount of flux carried by reaction/enzyme '$j$' is limited by the enzyme concentration ($E_j$) and the kinetic constants ($b_j$) which is based on enzyme turnover numbers[6], it can be represented as follows:

$$f_j = E_j \cdot b_j \qquad (7)$$

Now, substituting eq. (7) in (6), we get the additional flux constraint to be used which limits the sum of all intracellular fluxes within the enzyme capacities:

$$\sum_{j=1}^{N} a_j f_j \leq 1 \qquad (8)$$

where, $a_j = C \cdot v_j/b_j$, is referred to as 'crowding coefficient', C is cytoplasmic density which requires to be measured experimentally and $v_j$ is calculated using the molar mass ($M_j$) and specific volume of the protein (enzyme) as follows:

$$v_j = M_j \cdot v_j^{specific} \qquad (9)$$

Note that since the specific volume of the proteins ($v_j^{specific}$) is known to vary very minimally in different cells[17], we used the average value (0.73 mL/g) as reported earlier[6].

The additional constraint in FBAwMC, i.e. eq. (7), can be simply implemented in the S matrix itself. To do so, consider that stoichiometric matrix $S_{ij}$ and the flux vector $f_j$ are represented as below:

$$S_{ij} = \begin{bmatrix} s_{11} & s_{12} & \cdots & s_{1j} \\ \vdots & \vdots & \vdots & \vdots \\ s_{i1} & s_{i2} & \cdots & s_{ij} \end{bmatrix}$$

$$f_j = \begin{bmatrix} f_1 \\ \vdots \\ f_j \end{bmatrix}$$

Now, the additional constraint in FBAwMC based on crowding coefficients can be conveniently included in the S matrix in the form of a dummy reaction involving a pseudo metabolite whose sink flux is allowed to vary between 0 and 1 as follows.

$$S_{(i+1)(j+1)} = \begin{bmatrix} s_{11} & s_{12} & \cdots & s_{1(j+1)} \\ a_{1\ldots} & a_{2\ldots} & \cdots & a_{(j+1)\ldots} \\ s_{(i+1)1} & s_{i2} & \cdots & s_{(i+1)(j+1)} \end{bmatrix}$$

$$f_{j+1} = \begin{bmatrix} f_1 \\ f_{dr\ldots} \\ f_{j+1} \end{bmatrix}$$

where, $a_1, a_2 \ldots a_{(j+1)}$ are the coefficients of a single pseudo metabolite added to all reactions ranging from 1…. (j+1). Note that, $a_1, a_2 \ldots a_{(j+1)}$ are same as the crowding coefficients of reactions 1…. (j+1). The crowding coefficients of the dummy reaction, biomass objective, non-gene associated reactions and exchange reactions are set to zero. $f_{dr}$ represents the flux of the dummy reaction and is allowed to vary in the range of 0 and 1, satisfying enzyme capacity constraint.

#### *Estimation of cell crowding coefficients (a)*

The estimation of crowding coefficients required two kinds of experimental data: enzyme kinetic parameters, i.e. turnover numbers, $k_{cat}$ ($s^{-1}$) and cytoplasmic densities[6]. These are used in a specific manner to estimate the macromolecular crowding coefficients as explained in previous section. The enzyme turnover numbers were obtained from BRENDA database[18]. It should be noted that we used the turnover numbers specific to LAB whenever it is available. However, if it is not measured in LAB for any of the enzymes, then, the maximum reported value from any organism was used to avoid incorporation of smaller turnover numbers (higher crowding effects), which may limit the overall flux of the corresponding reactions. Cytoplasmic density of each LAB was estimated using the previously measured DCW values, volume occupied by a single cell and total number of cells in a given culture volume. Note that since cytoplasmic density is defined as the ratio of cell mass to cell volume, it was thus calculated after accounting for the necessary dilution factors.

### Supplementary Tables

#### Table S1. Genomic features of the six LAB examined in this study

| Species | Assembly | Genome size | GC% | # genes |
|---|---|---|---|---|
| *Lactobacillus plantarum* WCFS1 (LbPt) | GCA_000203855.3 | 3.34862 | 44.45 | 3174 |
| *Lactococcus lactis* subsp. *cremoris* NZ9000 (LcLt) | GCA_000192705.1 | 2.53029 | 35.7 | 2604 |
| *Lactobacillus casei* subsp. *casei* ATCC 393 (LbCs) | GCA_000829055.1 | 2.95296 | 47.86 | 2979 |
| *Lactobacillus salivarius* ATCC 11741 (LbSv) | GCA_000159395.1 | 2.01725 | 32.7 | 1983 |
| *Lactobacillus fermentum* ATCC 14931 (LbFm) | GCA_000159215.1 | 1.86701 | 52.6 | 1794 |
| *Leuconostoc mesenteroides* subsp. *mesenteroides* ATCC 8293 (LeMt) | GCA_000014445.1 | 2.07576 | 37.66 | 2089 |

#### Table S2. Characteristics of LAB genome-scale models

| Model properties | LbPt | LcLt | LbCs | LbSv | LbFm | LeMt |
|---|---|---|---|---|---|---|
| Genes (Unique) | 907 | 657 | 683 | 590 | 580 | 680 |
| Reactions | 1043 | 923 | 900 | 901 | 888 | 951 |
| - Metabolic | 861 | 769 | 725 | 743 | 732 | 795 |
| - Transport | 182 | 154 | 175 | 158 | 156 | 156 |
| - Gene associated | 952 | 834 | 802 | 817 | 800 | 868 |
| - Blocked reactions | 313 | 282 | 294 | 340 | 320 | 295 |
| - With $k_{cat}$ values | 498 | 468 | 445 | 442 | 427 | 484 |
| Metabolites | 950 | 897 | 880 | 868 | 879 | 900 |
| - Unique | 806 | 772 | 740 | 742 | 749 | 783 |
| - Dead ends | 94 | 238 | 81 | 90 | 86 | 83 |

**Table S3. OD600 conversion factors and cytoplasmic densities of 6 LAB**

| Species | Conversion factor (gDCW/L per OD600 unit) | Cytoplasmic density (gDCW/mL) |
|---|---|---|
| LbPt | 0.363 | 0.442 |
| LcLt | 0.405 | 0.623 |
| LbCs | 0.339 | 0.493 |
| LbSv | 0.393 | 0.637 |
| LbFm | 0.424 | 0.519 |
| LeMt | 0.494 | 0.388 |

**Table S4. Differential abundance of respective LAB in the corresponding probiotic-fed mice**

| Probiotic-fed mice condition | Corresponding LAB species | Log2FC wrt control | Adj. p-val |
|---|---|---|---|
| LbPt | LbPt | 3.4830 | 2.96e-5 |
| LcLt | LcLt | 0.7609 | 0.18593 |
| LbCs | LbCs | 7.6743 | 2.06e-6 |
| LbSv | LbSv | 8.8870 | 5.13e-8 |
| LbFm | LbFm | 4.895 | 0.0153 |
| LeMt | LeMt | 6.217 | 2.14e-7 |

### Supplementary Figures

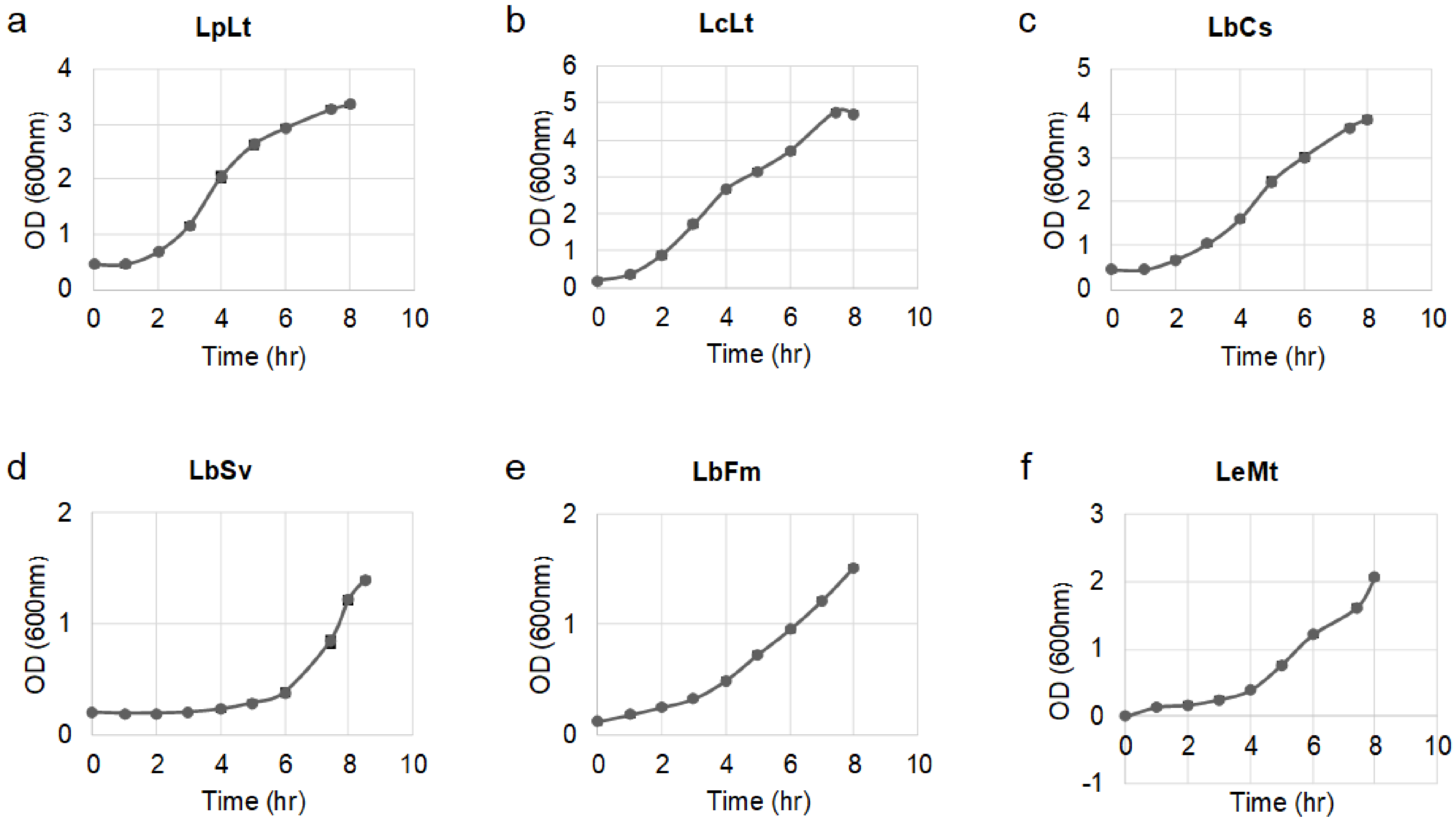

**Supplementary Figure S1. Growth characteristics of 6 LAB in LABDM.**

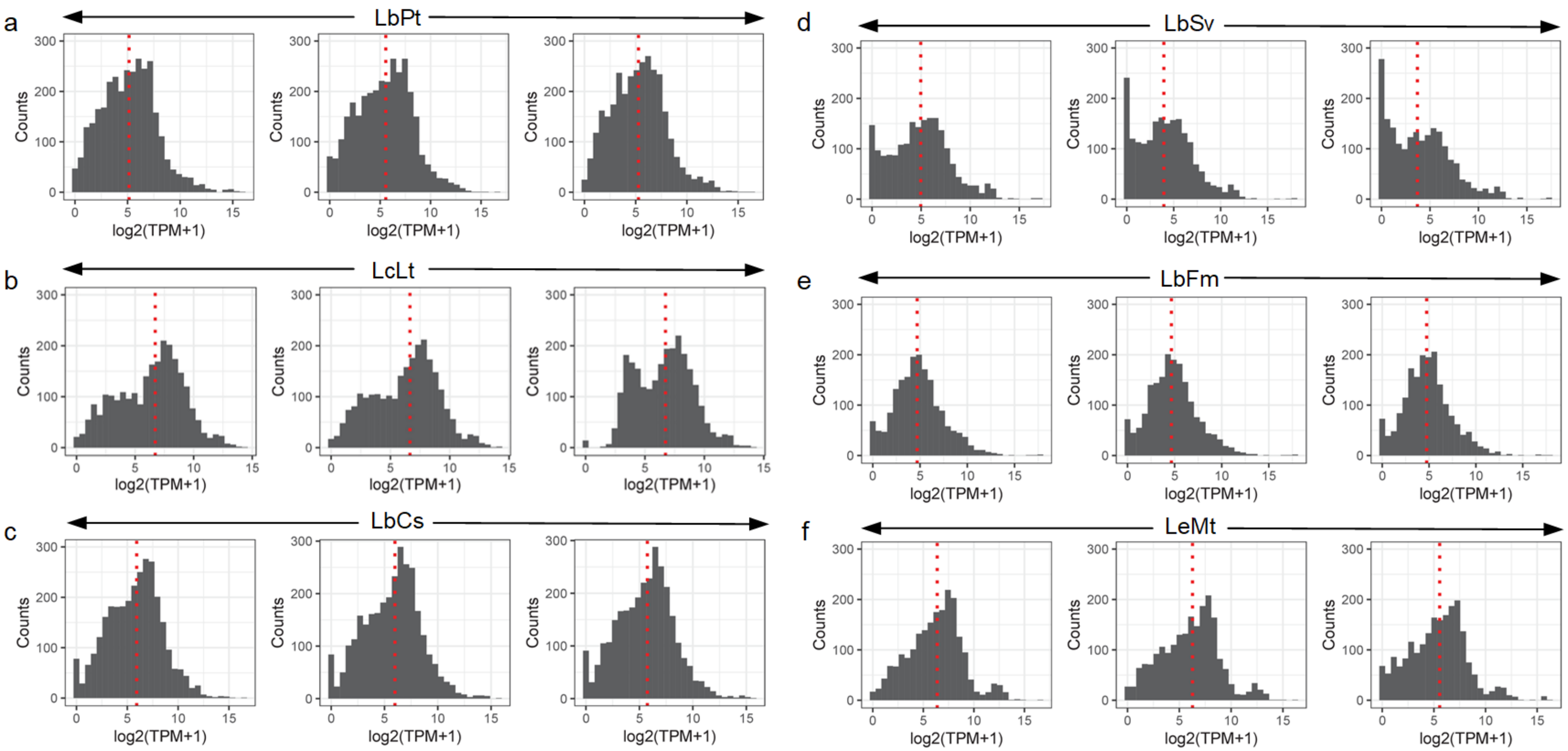

**Supplementary Figure S2. Genome-wide LAB transcriptome profiles.** Three plots in each LAB refer to the biological replicates and the vertical dotted red line indicate the median.

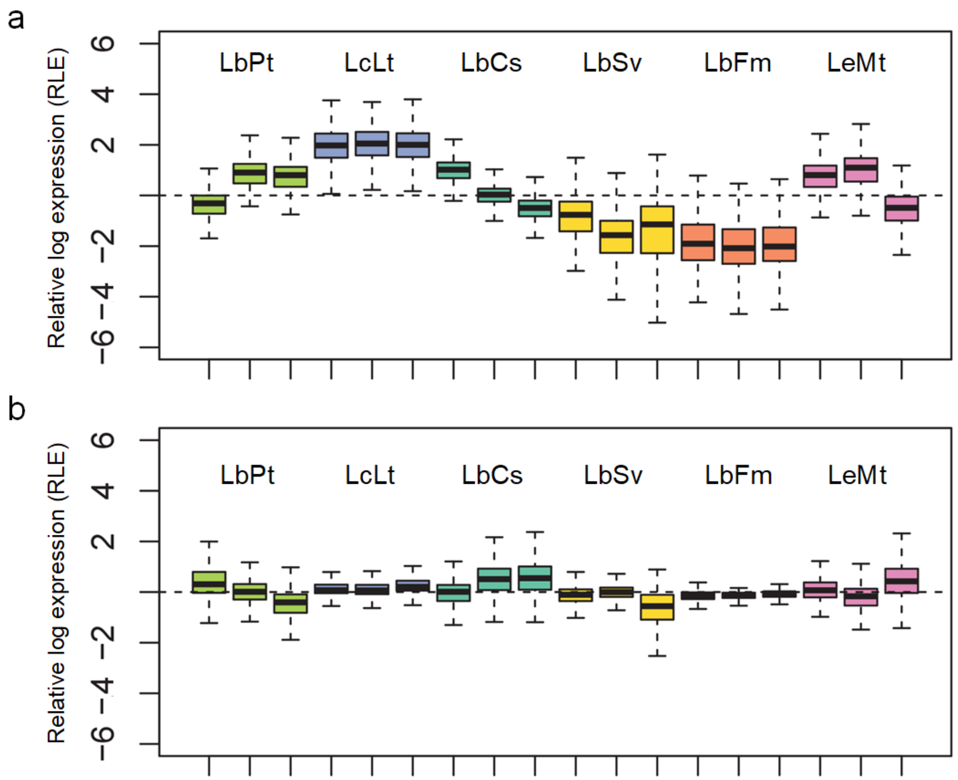

**Supplementary Figure S3. Relative log expression of orthologous genes before (a) and after (b) normalization w.r.t. to housekeeping genes.**

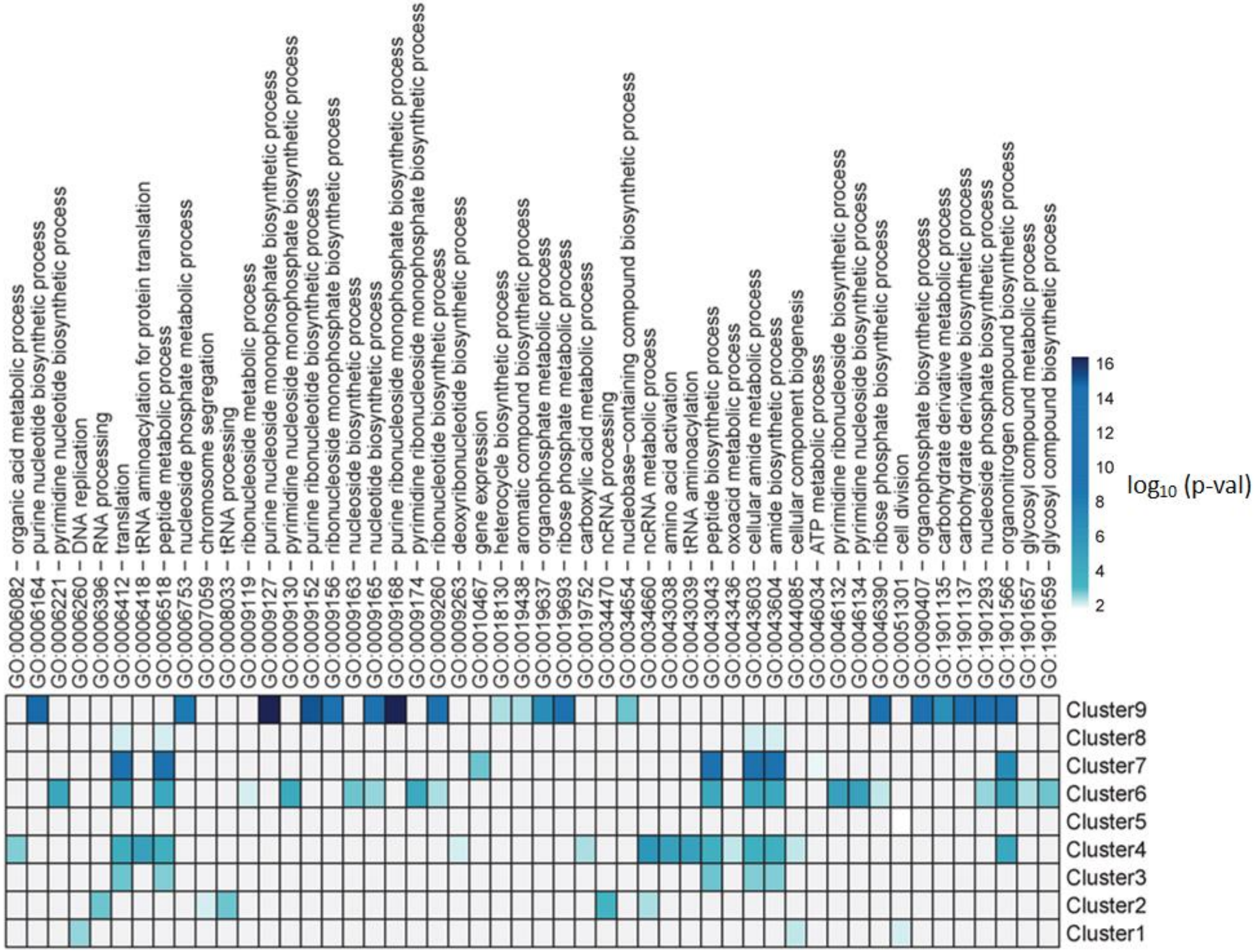

**Supplementary Figure S4. Enrichment of various GO categories in the various clusters of the orthologous gene presented in Figure 2.**

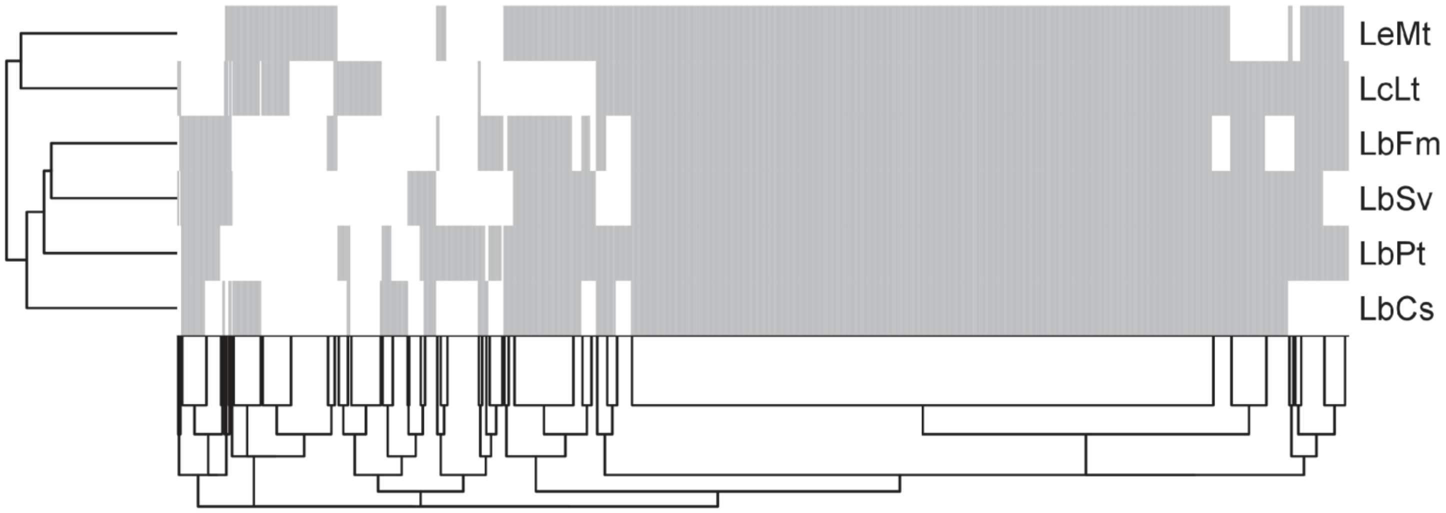

**Supplementary Figure S5. Hierarchical clustering of LAB based on reactome.** Euclidean distances were calculated for clustering from a matrix where the presence and absence of each reaction is represented in binary form, i.e. ‘0’ or ‘1’.

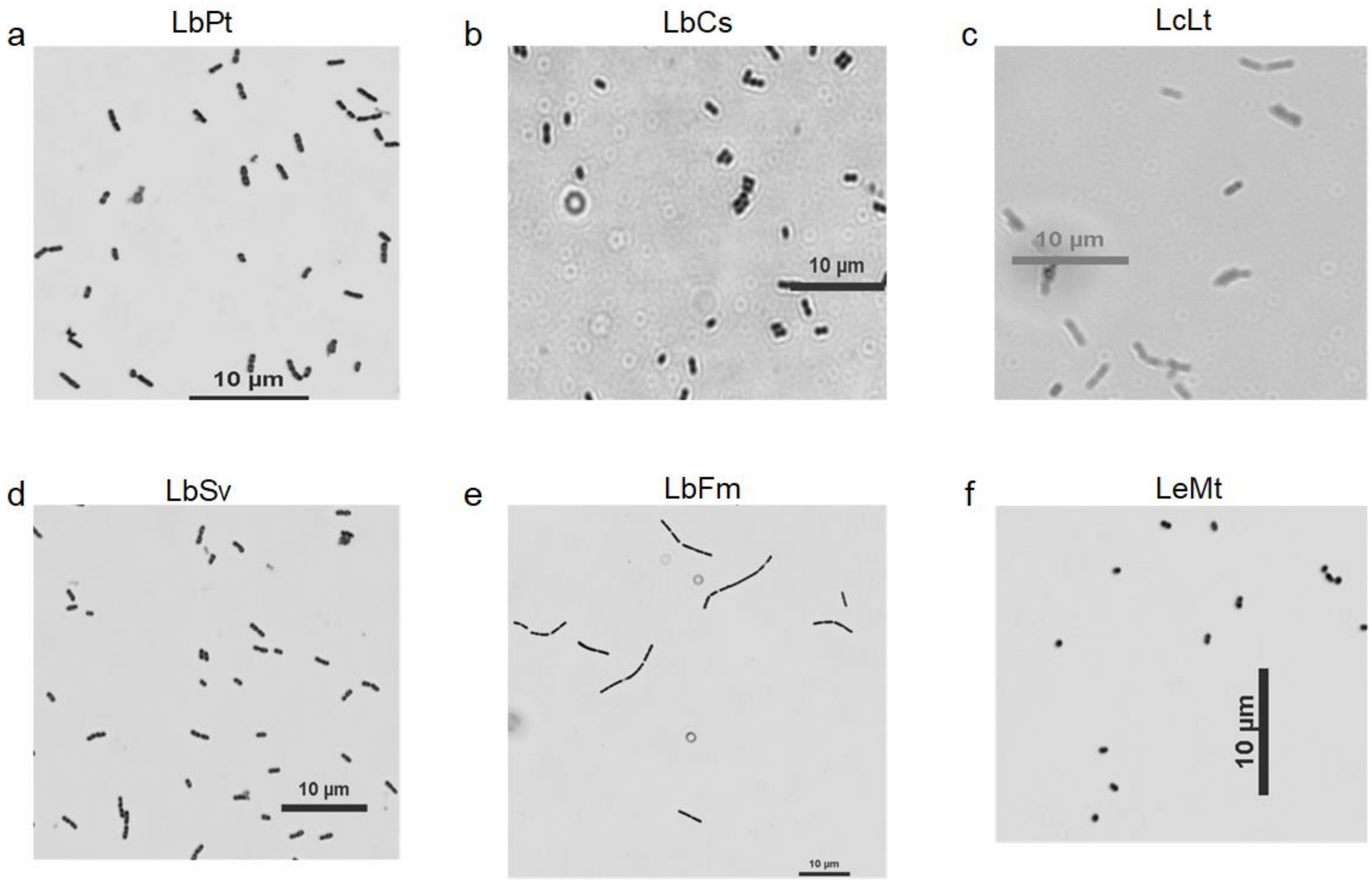

**Supplementary Figure S6. Cell sizes of LAB.** LAB sizes were determined using confocal microscopy.

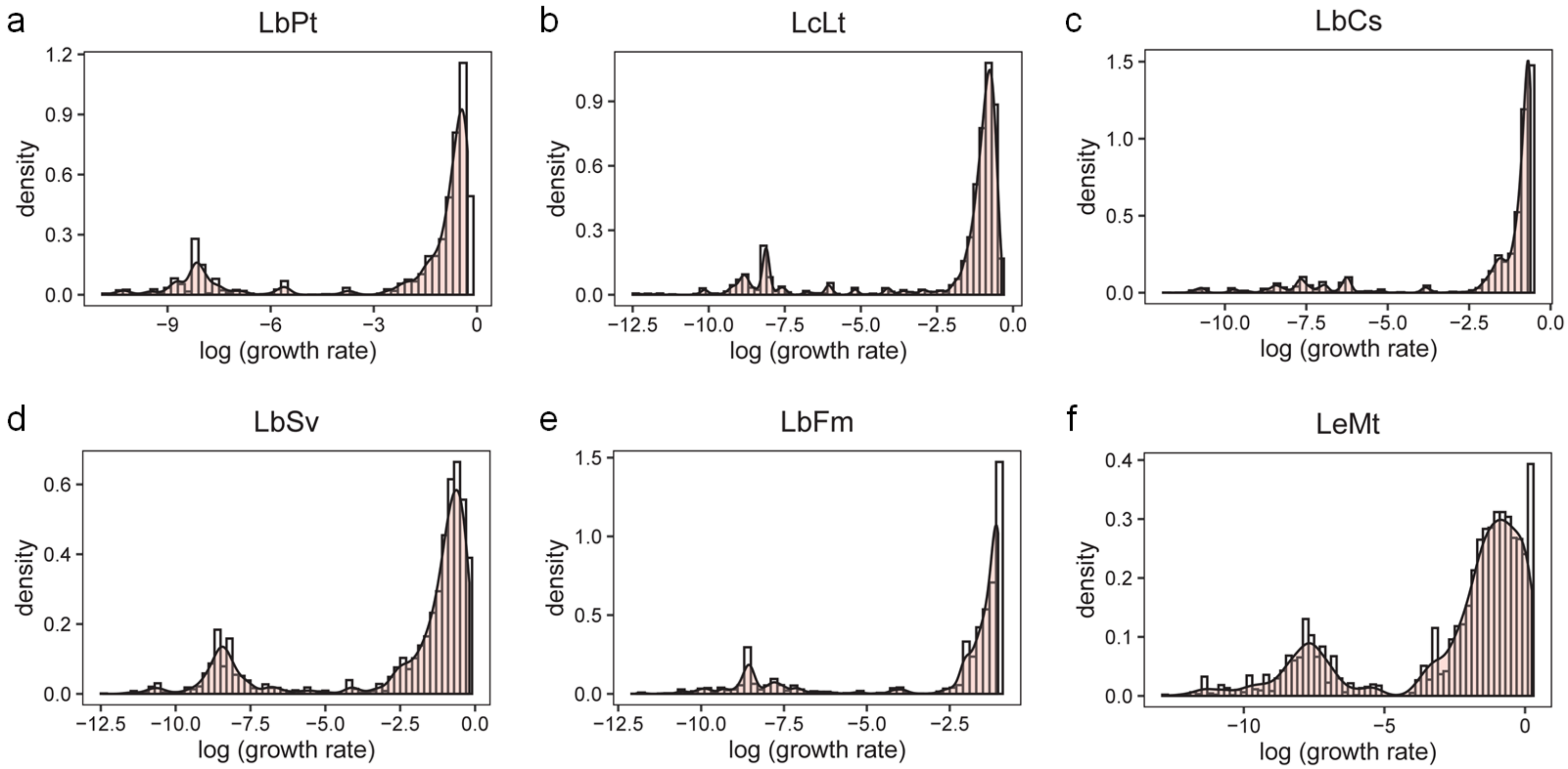

**Supplementary Figure S7. LAB growth characteristics as revealed by FBAwMC.** The distribution of log transformed growth rates from multiple $a_i$ parameters in 5000 samples is plotted.

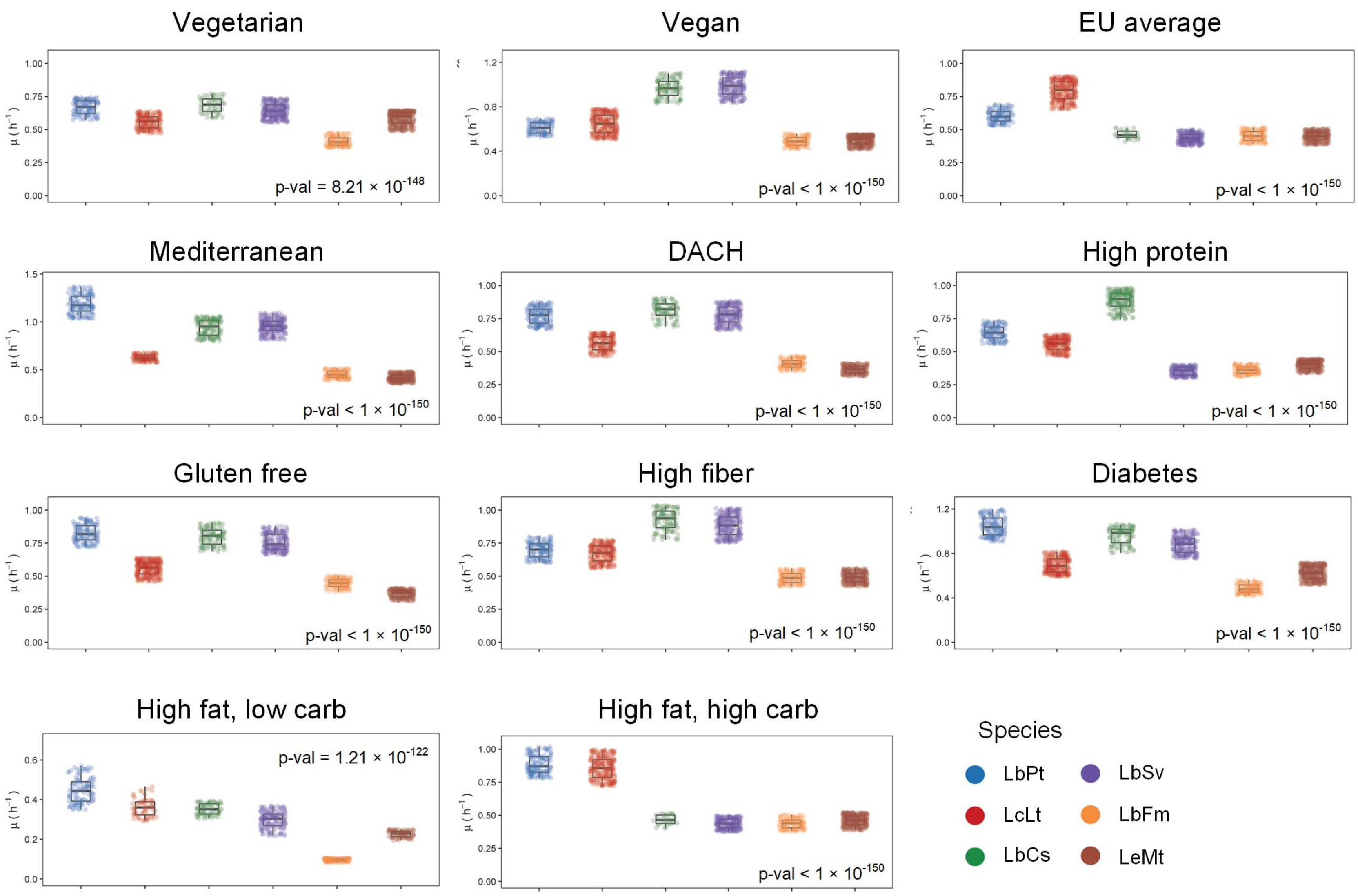

**Supplementary Figure S8. Variations in simulated growth rates across various diets in each LAB species** ($p$-val $< 1 \times 10^{-150}$, Kruskal-Wallis test)**.**

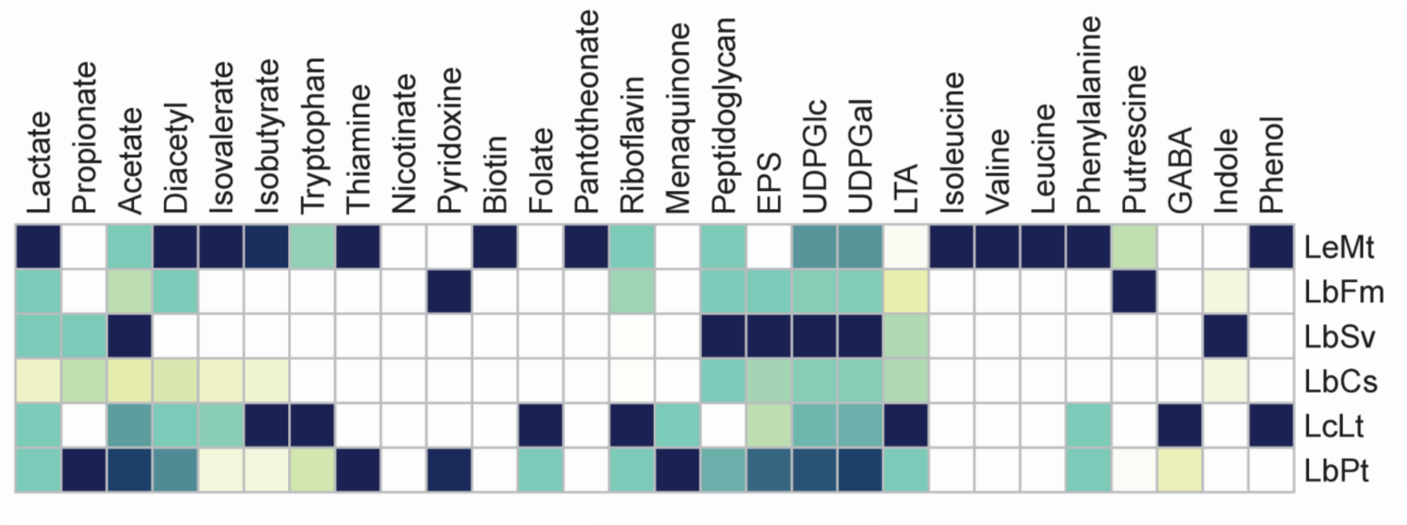

**Supplementary Figure S9. Postbiotic production capacities estimated using transcriptome constraints.**

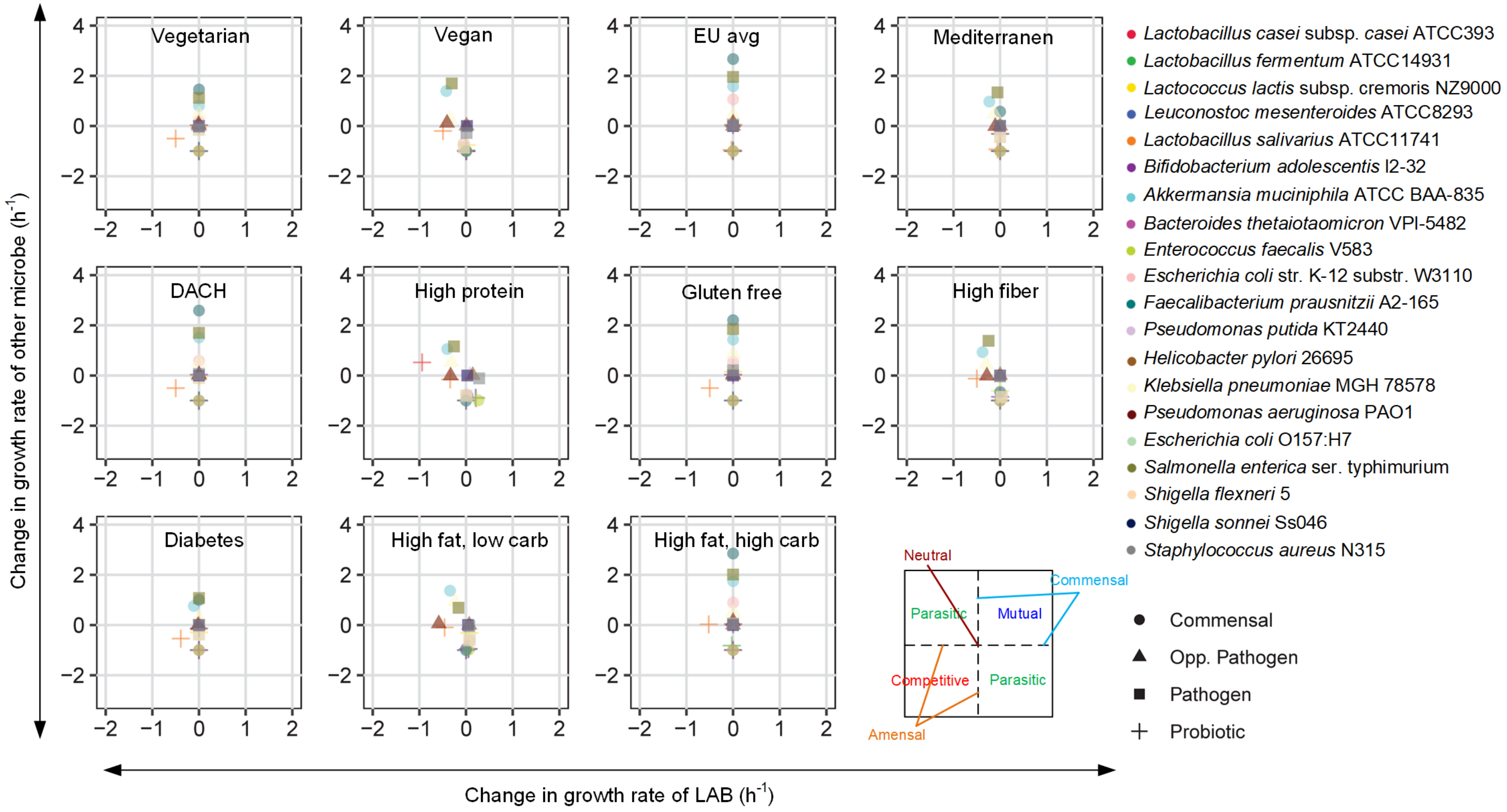

**Supplementary Figure S10. Interactions of LbPt with other gut microbes simulated *in silico*.** Outcomes of pairwise metabolic interactions with various other gut microbes. The relative difference in the growth rate of each strain when it grows separately or in combination with another organism is plotted.

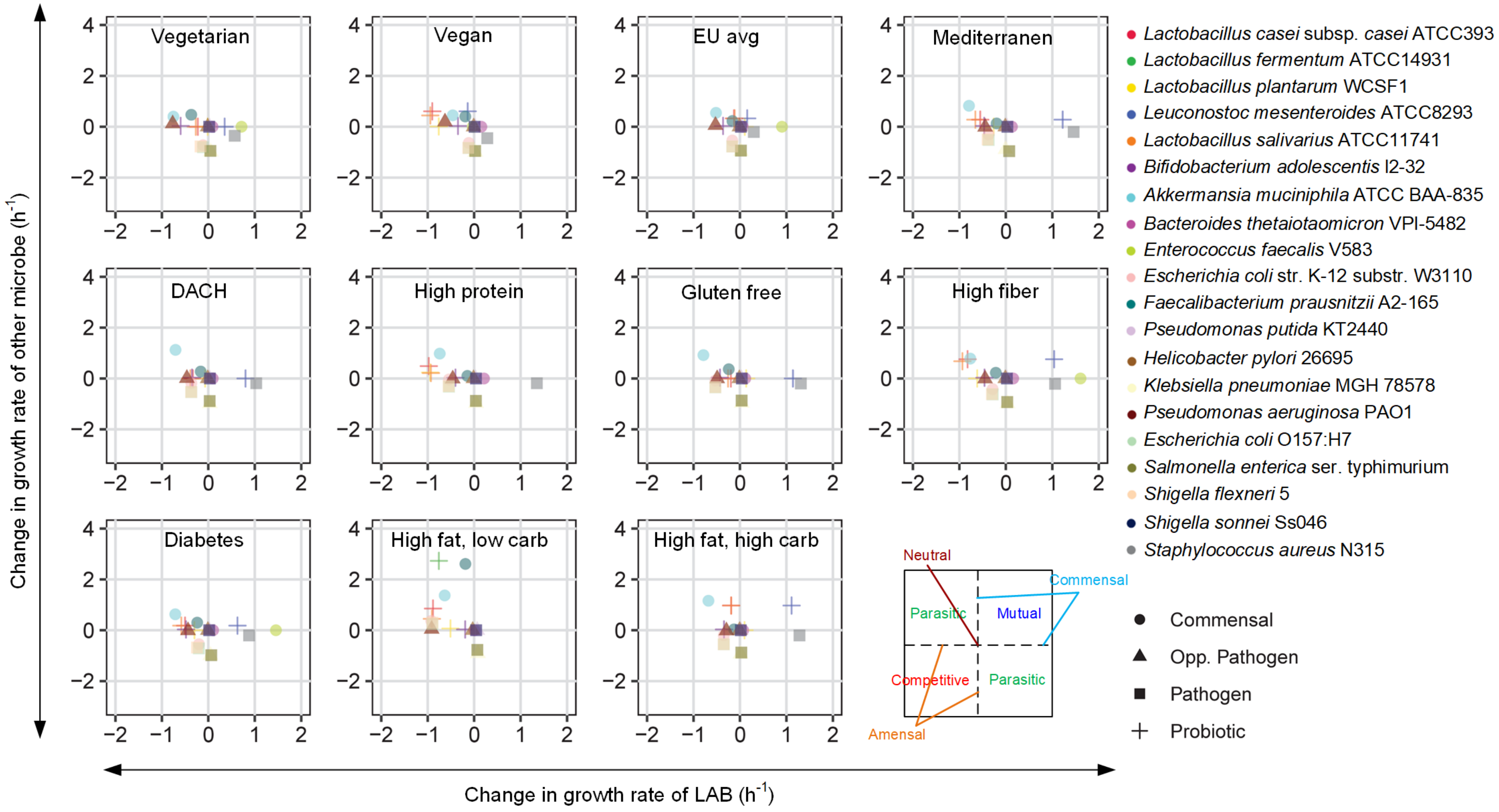

**Supplementary Figure S11. Interactions of LcLt with other gut microbes simulated in silico.** Outcomes of pairwise metabolic interactions with various other gut microbes. The relative difference in the growth rate of each strain when it grows separately or in combination with another organism is plotted.

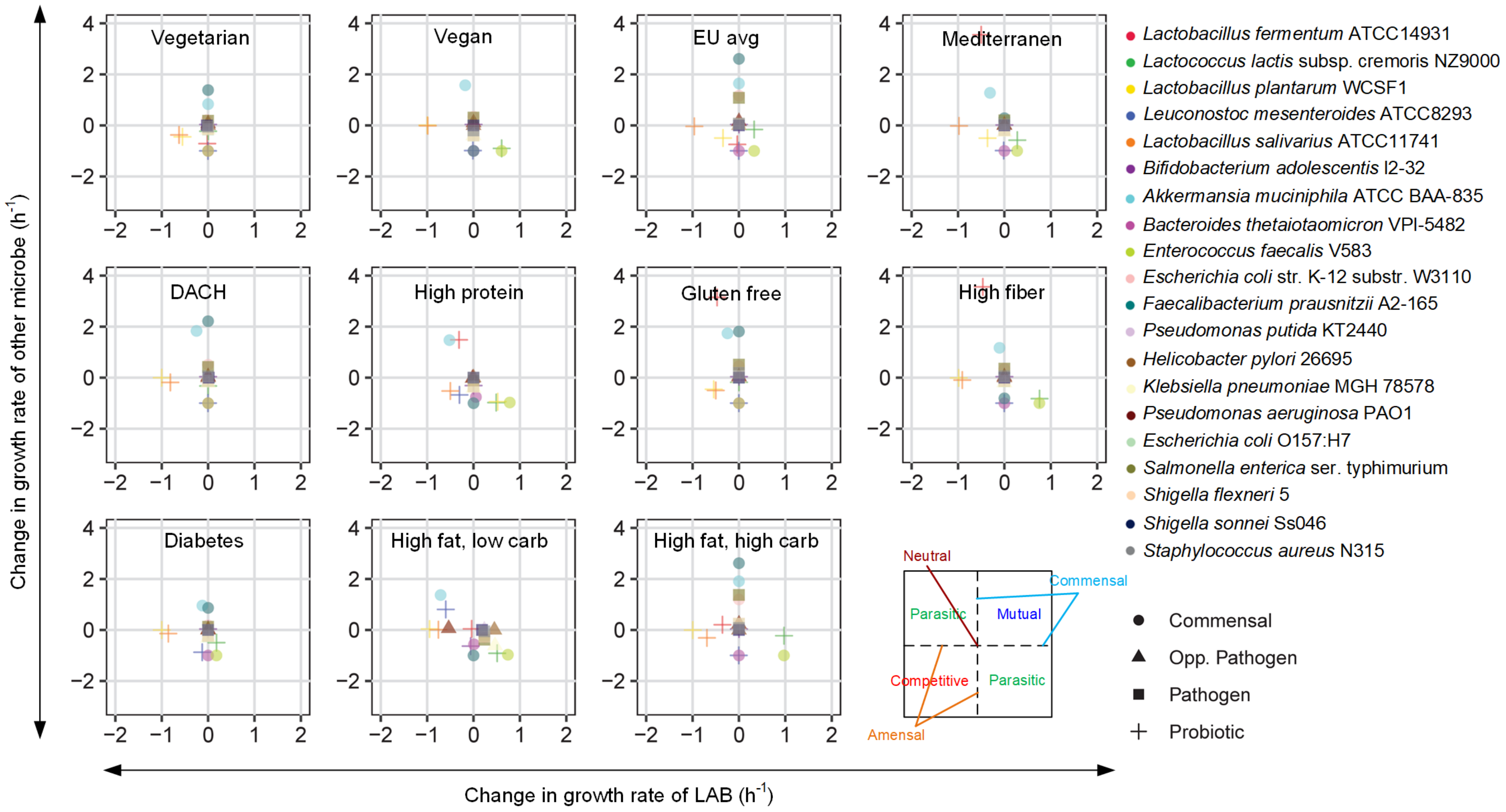

**Supplementary Figure S12. Interactions of LbCs with other gut microbes simulated in silico.** Outcomes of pairwise metabolic interactions with various other gut microbes. The relative difference in the growth rate of each strain when it grows separately or in combination with another organism is plotted.

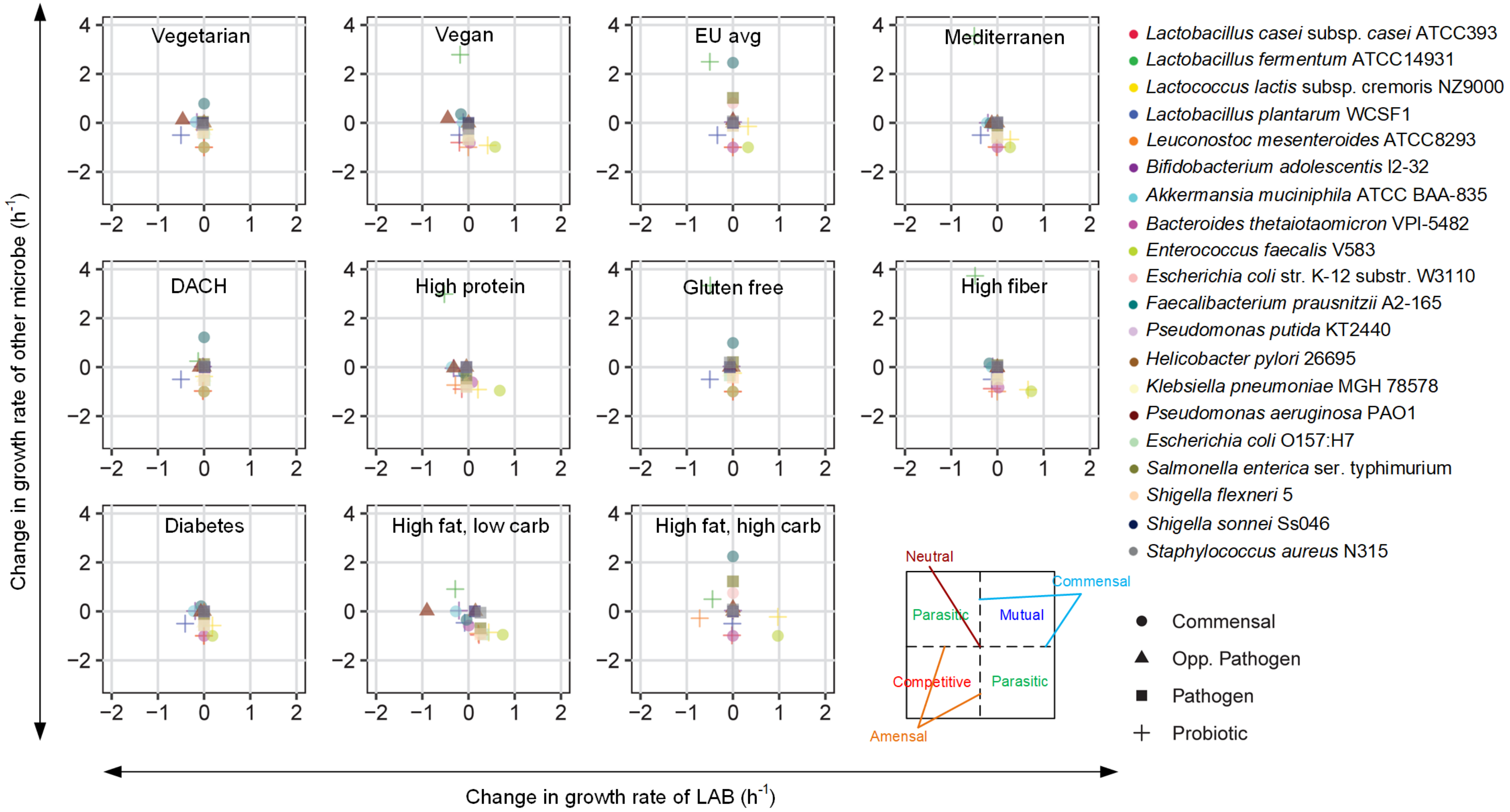

**Supplementary Figure S13. Interactions of LbSv with other gut microbes simulated in silico.** Outcomes of pairwise metabolic interactions with various other gut microbes. The relative difference in the growth rate of each strain when it grows separately or in combination with another organism is plotted.

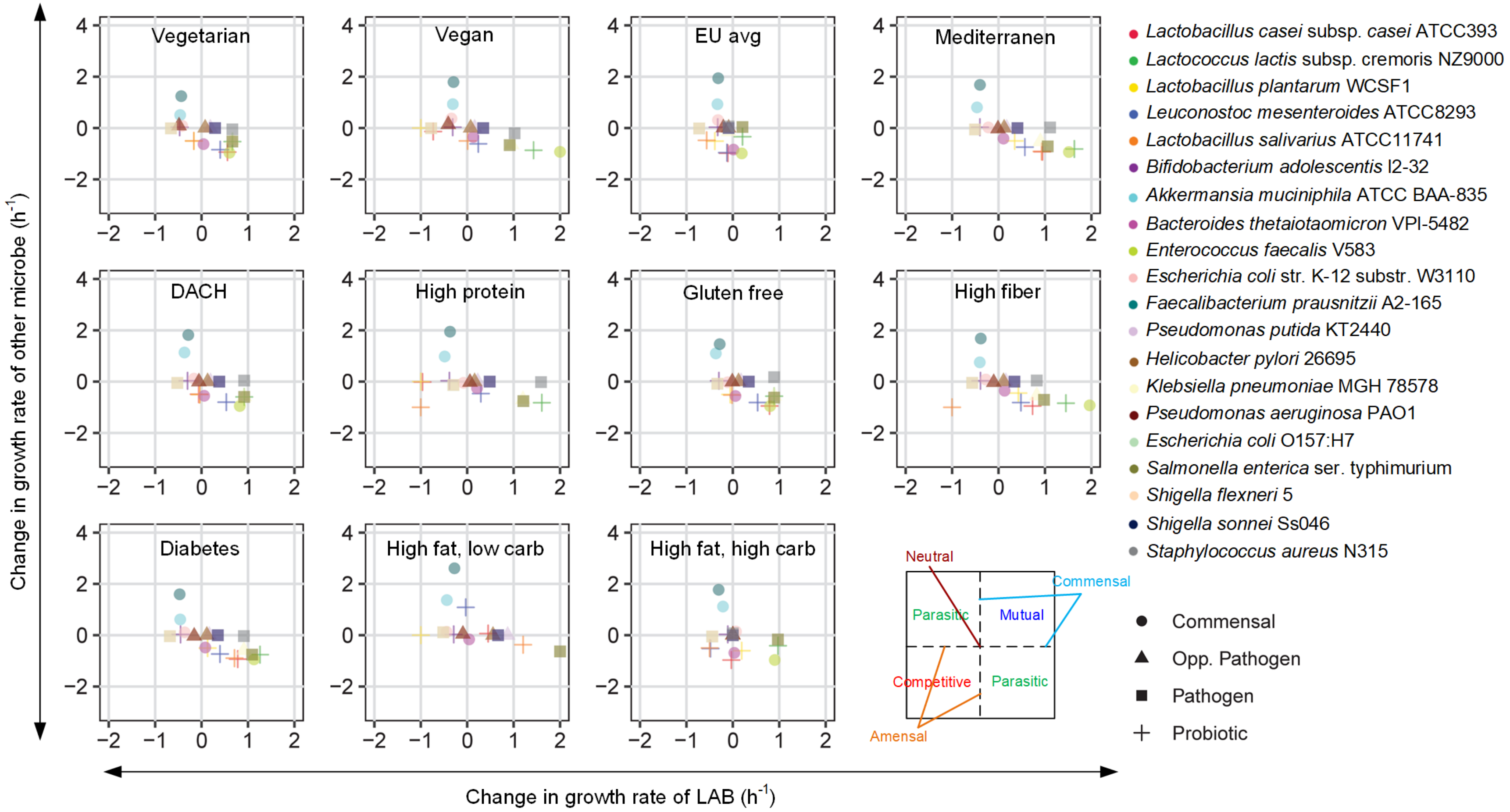

**Supplementary Figure S14. Interactions of LbFm with other gut microbes simulated in silico.** Outcomes of pairwise metabolic interactions with various other gut microbes. The relative difference in the growth rate of each strain when it grows separately or in combination with another organism is plotted.

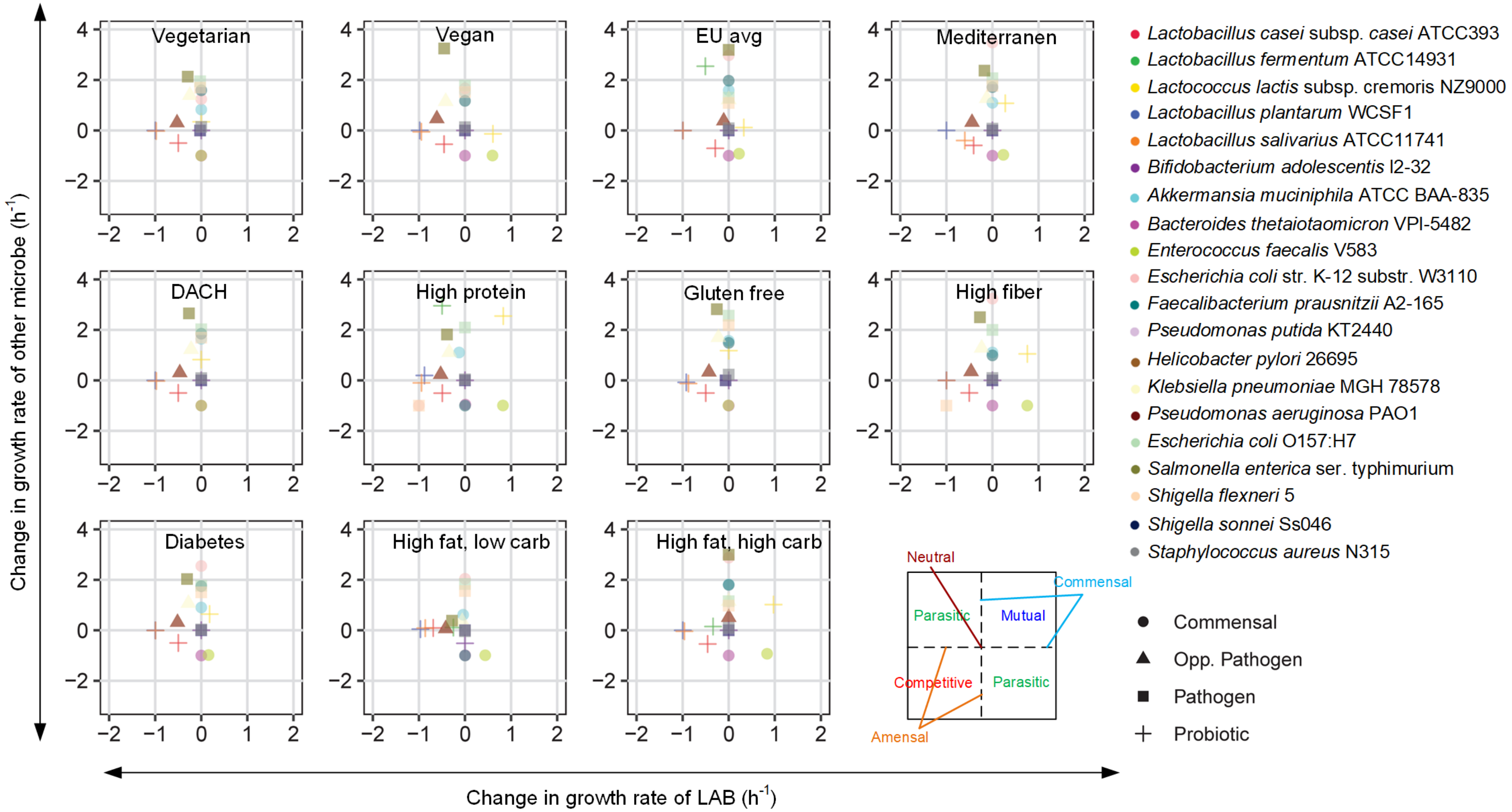

**Supplementary Figure S15. Interactions of LeMt with other gut microbes simulated in silico.** Outcomes of pairwise metabolic interactions with various other gut microbes. The relative difference in the growth rate of each strain when it grows separately or in combination with another organism is plotted.

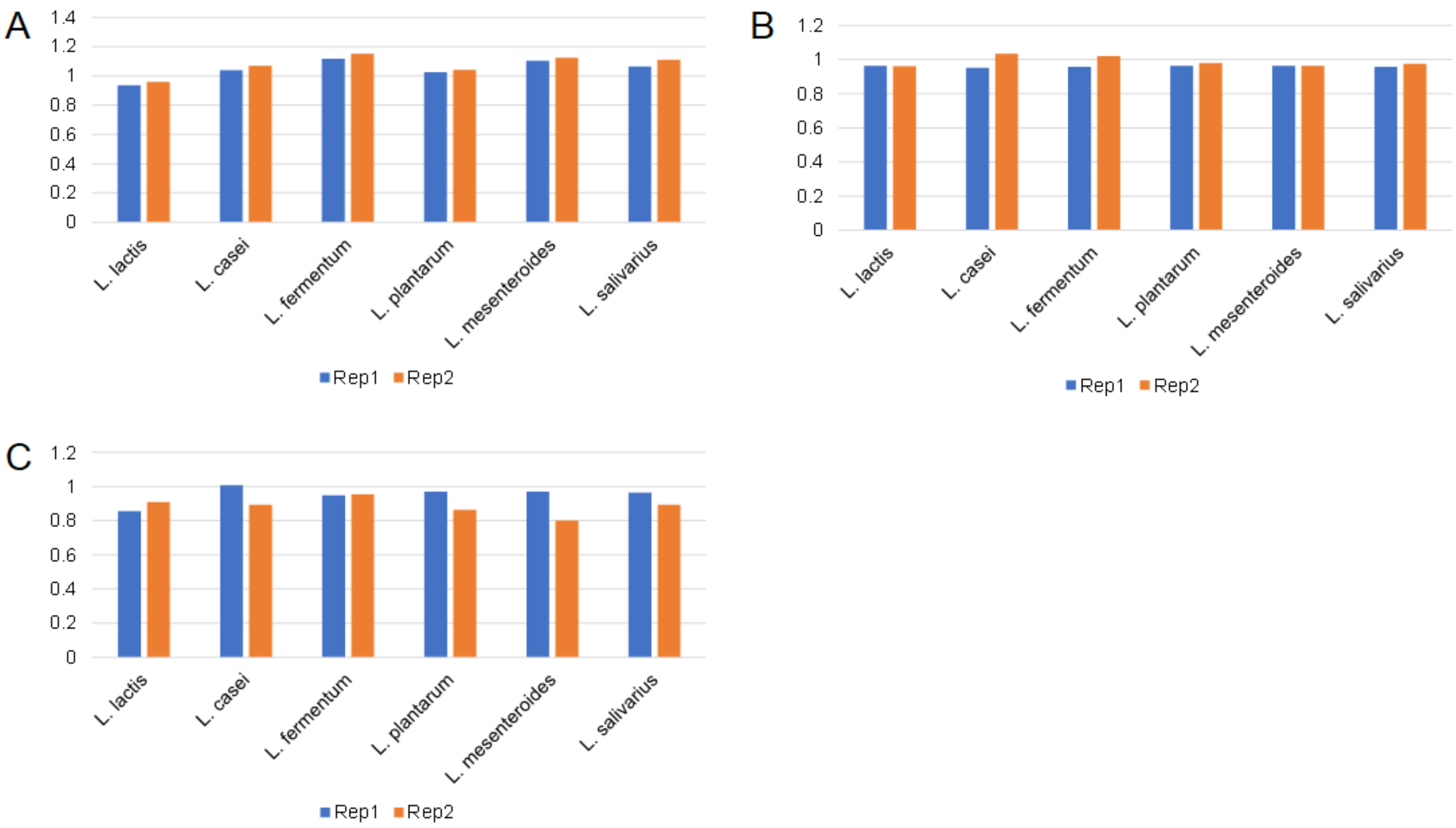

**Supplementary Figure S16. Growth of commensals in spent-medium transfer experiments.** Fold-change of commensal species' growth with and without LAB supernatants is reported. (A) *A. muciniphila*, (B) *B. thetaiotamicron* and (C) *E. coli*.
